# miRNA-29-CLIP uncovers new targets and functions to improve skin repair

**DOI:** 10.1101/2024.05.02.591622

**Authors:** Lalitha Thiagarajan, Rosa Sanchez-Alvarez, Chiho Kambara, Poojitha Rajasekar, Yuluang Wang, François Halloy, Jonathan Hall, Hans-Jürgen Stark, Iris Martin, Petra Boukamp, Svitlana Kurinna

## Abstract

MicroRNAs (miRNAs) control organogenesis in mammals but their role in specific cell types is not fully explored. miRNAs exert their function by binding mRNAs and inhibiting translation. Skin is an excellent model to study the role of miRNAs in primary cells of epidermal (keratinocytes) and mesodermal (fibroblasts) origin, because the growth of these cells is tightly controlled at translation. Previous research demonstrated that miRNA-29 family functions during skin repair, however, the exact mRNA targets and the downstream mechanisms of miRNA-29-mediated regulation of cell growth is missing. Here, we use miRNA crosslinking and immunoprecipitation (miRNA-CLIP) method to find the direct targets of miRNA-29 in keratinocytes and fibroblasts from human skin. We uncover previously unrecognized roles of miRNA-29 in protein folding and RNA processing, common to all cell types tested, and determine the cell-specific role of miRNA-29. Using modified anti-sense oligonucleotides (ASO) in 2D and 3D cultures of keratinocytes and fibroblasts, we enhanced cell-to-matrix adhesion and found an autocrine and paracrine mechanism of miRNA-29-dependent cell growth. Our results include a comprehensive list and functional analyses of mRNAs directly bound by miRNA-29 keratinocytes and fibroblasts, determined by miRNA-CLIP and ASO-mediated inhibition of miRNA-29 followed by RNA-seq. We reveal a full transcriptome of human keratinocytes with enhanced adhesion to the wound matrix, which supports regeneration of the epidermis and is regulated by miRNA-29. The functions of miRNA-29 identified in this study can provide a new approach to improve cutaneous repair by restoring and enhancing the endogenous mechanisms through the stage-specific delivery of miRNA-29 ASO.

## INTRODUCTION

MicroRNAs (miRNAs) are short non-coding RNAs that regulate distinct functions in mammalian cells by direct binding and suppression of mRNAs. miRNAs regulate cell plasticity and are especially important in defining cell fate during differentiation and regeneration of tissues and organs. miRNA crosslinking and immunoprecipitation (miRNA-CLIP) can identify miRNA targets in a cell-type specific manner by directly cross-linking and precipitating functional AGO2 complexes of miRNA-mRNA bound by the ‘seed’-sequence match^1–3^. If coupled with the gain- and loss-of-function experiments, this method has the potential to elicit the function of a miRNA in the most direct, functional way, identifying both new miRNA-mRNA pairs and placing the known mRNA targets into biological pathways regulated by the miRNA.

Skin regeneration is an excellent model of adult tissue- and organogenesis, which is tightly controlled at both transcriptional and translational level by epigenetic factors, including miRNAs^4^. Recent advances in treating severe skin conditions with autologous transgenic therapies emphasize the importance of modulating the molecular mechanisms involved in repair^5^. However, the mechanisms of miRNAs activity toward cell-specific transcriptomes in the skin epidermis and in the underlying dermis are largely missing. *In silico* and *in vitro* analyses can only partially identify the *in vivo* functions of miRNAs, because these functions depend on the expression of miRNA-RNA pairs in the epidermal and dermal cells during the specific stage of their cell cycle or differentiation. We demonstrated that miRNA-CLIP allows unbiased identification of mRNA targets in epidermal cells (keratinocytes) and in dermal fibroblasts using RNA sequencing as a read-out, which we then combined with the functional assays, specific for keratinocytes and fibroblasts. As an example, we successfully implemented this approach to unveil the complex targetome of miRNA-29 family of miRNA-29a/b/c, which are known to function in developing normal cells^6–8^ and in cancer^9^. In mammals, the miRNA-29 family is encoded by two genes carrying four precursor sequences, pre-miRNA-29a, pre-miRNA-29b1, pre-miRNA-29b2, and pre-miRNA-29c, processed into three functional miRNA-29a-3p, miRNA-29b-3p, and miRNA-29c-3p with a common ‘seed’ sequence. We and others previously detected very little endogenous activity for miRNA-29b^6, 10, 11^, suggesting that most of miRNA-29 functions are exerted by miRNA-29a and miRNA-29c, which differ by only one nucleotide outside of the ‘seed’ region. An unusually versatile targetome of the miRNA-29 family includes mRNAs that regulate cell cycle ^12^, fibrosis ^13^, apoptosis^14^, and cell differentiation^15^. Our published and unpublished results supported by the reports on miRNA-29 functions in different organs and tissues have suggested a role for miRNA-29 in regenerating keratinocytes ^6, 8, 10^ and in dermal fibroblasts^13, 16^. Here, we used human skin as a model to unveil the cell-specific miRNA-29 targetome of keratinocytes and fibroblasts during their growth and differentiation in 2D and 3D models of human skin *ex vivo*. We tested the psoralen-biotin-modified precursor of miRNA-29, which was processed by the Dicer into an active miRNA-29 and incorporated into to a functional RISC. We then proceeded with the miRNA-29-CLIP experiments in primary human keratinocytes and fibroblasts, which revealed a function of miRNA-29 in cell adhesion and extracellular matrix (ECM). We confirmed this by identifying specific regulators of rapid keratinocyte adhesion, a function essential for clonal expansion in regenerative medicine of the skin^5^. Altogether, we reported the transcriptome directly regulated by miRNA-29 in human hair follicle, interfollicular keratinocytes, and in dermal fibroblasts. Using antisense miRNA-29 oligonucleotides (ASOs), we also discovered a new endogenous function of miRNA-29 in basal keratinocyte adhesion, fibroblast proliferation, and the formation of the ECM, which in turn, enhanced the rapid adhesion of keratinocytes through paracrine mechanisms. We identified FERMT2, SPARC, and COL4 as master regulators of keratinocyte adhesion and ECM deposition downstream of miRNA-29. Our results demonstrated that inhibition of miRNA-29 in keratinocytes and fibroblasts can be used to improve skin regeneration.

## RESULTS

### miRNA-29 defines epidermal growth in mouse wounds and in human skin equivalents

The growth and formation of the epidermis during development and regeneration of the skin implicates strong and rapid adhesion of the epidermal cells (keratinocytes) to basal lamina, continuous proliferation and differentiation of keratinocytes into the subrabasal layers^17^. Small excisional wounds created on mouse dorsum closed within a week, regenerating epidermis very rapidly through proliferation and migration of the surrounding intact epidermis. Inside the wound, the newly formed epidermis has more layers compared to an unwounded skin and in this, resembles human epidermis. Using *in situ* hybridization, we detected a strong signal for miRNA-29a in all layers of mouse epidermis except the lowest single-cell layer of keratinocytes attached to basal lamina (Figure 1). Epidermis in the middle of the wound where basal keratinocytes migrated and adhered to fibronectin did not show detectable levels of microRNA-29, with its expression activated in imidiate suprabasal keratinocytes (Figure 1A). In light of the known function of miRNA-29 in suppressing strong desmosomal adhesion in mouse and human epidermis^8^, we hypothesized that the molecules establishing basal adhesion in wounds has to be releaved of miRNA-29-mediate repression to help keratinocyte migrate and attach to provisional matrix in the dermis. These processes are crucial during wound closure as they regulate balanced migration and proliferation of keratinocytes, avoiding a chronic wound scenario. This balance may well be regulated by different levels of miRNA-29 as it can suppress adhesion to allow both lateral and longitudinal migration of keratinocytes in the wound. We thus decided to investigate the mechanism of miRNA-mediated adhesion of newly forming epidermis as it attached to the underlying dermis. We observed more signal for miRNA-29 in suprabasal epidermis in the peripheral part of the wound as well as in the unwounded epidermis (Figure 1B). In human skin equivalents (SE), overexpression of microRNA-29a/b mimic in keratinocytes seeded on dermal scaffolds of growth-arrested fibroblasts and the ECM^18^, resulted in suprabasal marker keratin K10 prematurely expressed in the basal layer, while the SE with non-specific mimic control had K10-negative basal layer (Figure 1C). Overexpression of miRNA-29 also resulted in thinner epidermis (Suppl. Figure S1A) suggesting loss of basal adhesion and premature differentiation. These co-cultures of keratinocytes and fibroblasts in 3D, also known as organotypic skin cultures^19^, resemble growth and stratification of human epidermis when exposed to the air-liquid interphase and can be grafted onto open wounds in rodents^20^ and in human patients^5^. Our data suggested that inhibition of miRNA-29 could improve basal adhesion and growth of SE, a desirable outcome for tissue regeneration and engineering.

**Figure 1.**
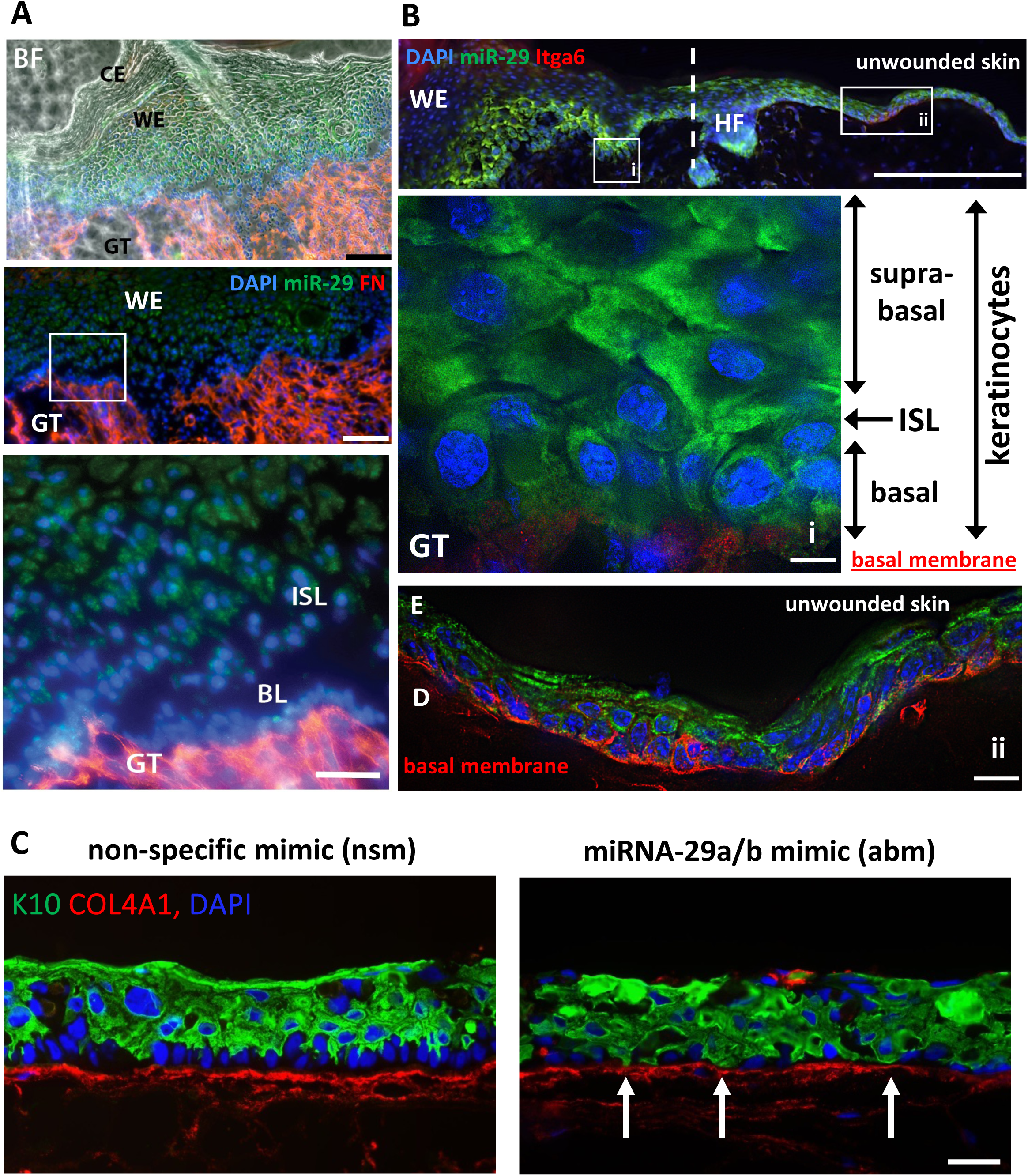
miRNA-29 is differentially expressed in regenerating mouse epidermis and induces premature differentiation of keratinocytes in human skin equivalents. (A) Fully restored epidermis of a closed cutaneous wound at day 5 of healing shows miRNA-29 in keratinocytes. Bright field (BF) shows cornified envelope (CE) on newly formed wound epidermis (WE) with high levels of miRNA-29, whereas the immediate suprabasal layer of the epidermis (ISL) and especially the basal layer (BL) attached to the granulation tissue (GT) express very little miRNA-29. miRNA-29 is detected by in situ hybridization *(green)* co-stained with fibronectin (red), and DAPI (blue). Scale bar = 100μm and 50μm. (B) Away from the wound bed, miRNA-29 is expressed mainly in the suprabasal epidermis with the evidence of polarized expression in keratinocytes transitioning from basal to differentiated suprabasal state, shown in (i). miRNA-29 is *green*, nuclei are *blue*. Integrin alpha 6 (Itga6, *red*) shows adhesion to the basal membrane connecting epidermis to the underlying dermis. WE – wound epidermis, GT – granulation tissue, (ii) - unwounded skin, with intact E – epidermis, D – dermis. HF is a hair follicle, together with punctate line marking the arbitrary border between wounded skin and the adjacent normal skin. Scale bars are 200μm, 10μm, and 20μm. (C) Human skin equivalents (SE) grown in 3D cultures at air-liquid interface for 6 days, stained with the markers of early differentiation keratin K10. The border between epidermis and the dermal scaffold is stained with collagen IV (COL4A1). Arrows indicate early expression of keratin K10 in the basal layer of keratinocytes transfected with miRNA-29a/b mimic (abm), whereas in the SE transfected with a non-specific mimic (nsm) keratin K10 expression starts at the immediate suprabasal layer and doesn’t “touch” collagen IV. Scale bar = 50μm.

Indeed, keratinocytes transfected with the single-stranded complimentary oligonuclotide inhibitor (ASO) of all miRNA-29a/b/c (abc), formed SE just as well as controls (non-specific ASO, nsa) while miRNA-29 mimic-transfected SE struggled to attach through the integrin beta1 (ITGB1)-mediated adhesion (Figure 2A). A major adhesion receptor and progenitor marker in the skin^21^, ITGB1 showed a better deposition and distribution in SE treated with miRNA-29 ASO (Figure 2A). ITGA6, a ligand of laminin 332 responsible for basal adhesion of keratinocytes, was also deposited more in abc-SE to ensure epidermis-to-dermis adherence, marked by collagen 4 (Figure 2A). Importantly, BrdU incorporation rate did not change in response to miRNA-29 expression or its knock-down (Figure 2B, quantified in Suppl. Figure S1C), confirming that miRNA-29 does not regulate proliferation of keratinocytes.

**Figure 2.**
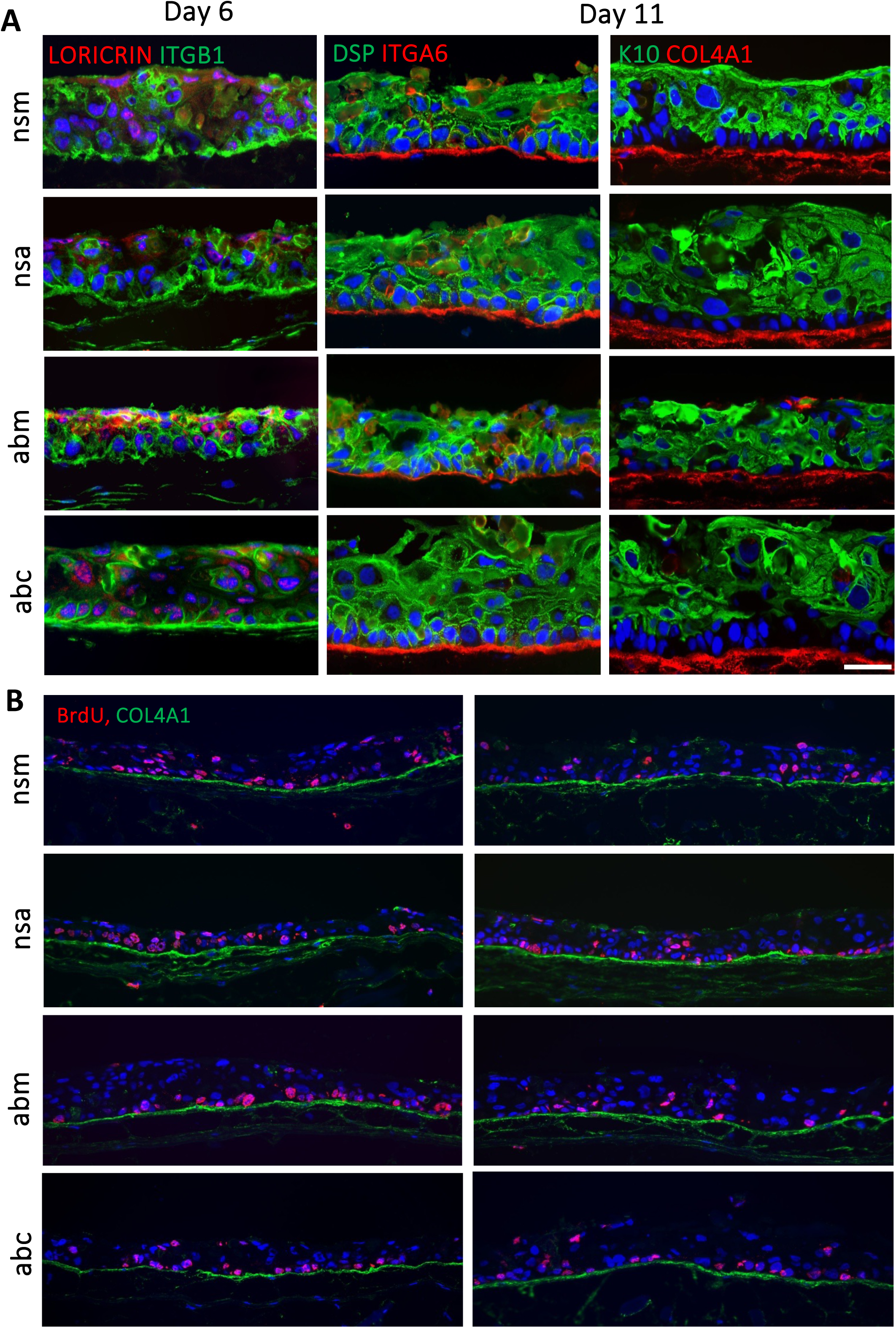
Inhibition of miRNA-29 supports growth of keratinocytes in human skin equivalents. **(A)** Human skin equivalents (SE) grown in 3D cultures at air-liquid interface for 6 and 11 days, and stained with the markers of basal adhesion (ITGB1 and ITGA6), early (keratin K10) and late differentiation (LORICRIN), and cell-to-cell adhesion (desmoplakin DSP). The border between epidermis and the dermal scaffold is stained with collagen IV (COL4A1). Scale bar = 50μm. **(B)** BrdU staining of SE sections of keratinocytes transfected with non-specific mimic (nsm), non-specific inhibitor (nsa), miR-29 mimic (abm), miR-29 inhibitor (abc). Scale bar = 100μm.

These results demonstrate that miRNA-29 expression correlates with the loss of basal adhesion *in vivo*, which may be important for the acquisition of suprabasal barrier-forming functions by regenerating epidermis. Over-expression of miRNA-29 inhibited growth of human epidermis, whereas suppression of miRNA-29 supported growth of keratinocytes possibly, by improving their basal adhesion.

### Rescue of miRNA-29 mRNA targets improves basal adhesion of human keratinocytes

It is well established that rapid adhesion of primary human keratinocytes is essential for their proliferation and expansion^5^. To test if low levels of miRNA-29 improve basal adhesion, we used a rapid adhesion assay following overexpression or inhibition of miRNA-29. High levels of miRNA-29 abrogated the rapid adhesion of keratinocytes and stopped their further proliferation, whereas loss of miRNA-29 significantly increased the number of rapidly attaching cells (Figure 3A, B, Suppl. Figure S2). Loss of miRNA-29 allowed cells to form clusters, essential for keratinocyte growth and maintenance of regenerative potential (Figure 3A). Importantly, enhanced adhesion following inhibition of miRNA-29 was not dependent on the presence of growth factors in the cell culture medium (Figure 3C), an observation important for potential therapeutic applications. Consistently with our previous data in human SE, loss of miRNA-29 did not increase keratinocyte proliferation as may be suggested by a higher number of cells in the adhesion assay (Suppl. Figure 1C). Moreover, when miRNA-29 inhibition experiment went beyond day 4 and until day 8, the control cells became cobble-like and showed signs of spontaneous differentiation expected from primary keratinocytes growing in 2D culture for 7 days^6^. In contrast, cells with loss of miRNA-29 function survived multiple transfections and continued proliferating until day 6, showing morphology of expanding primary cells in 2D culture until day 8 (Suppl. Figure 3). This is an important finding suggesting that inhibition of miRNA-29 could be used to support the expansion of primary keratinocytes, and that it would happen primarily through improved adhesion.

**Figure 3.**
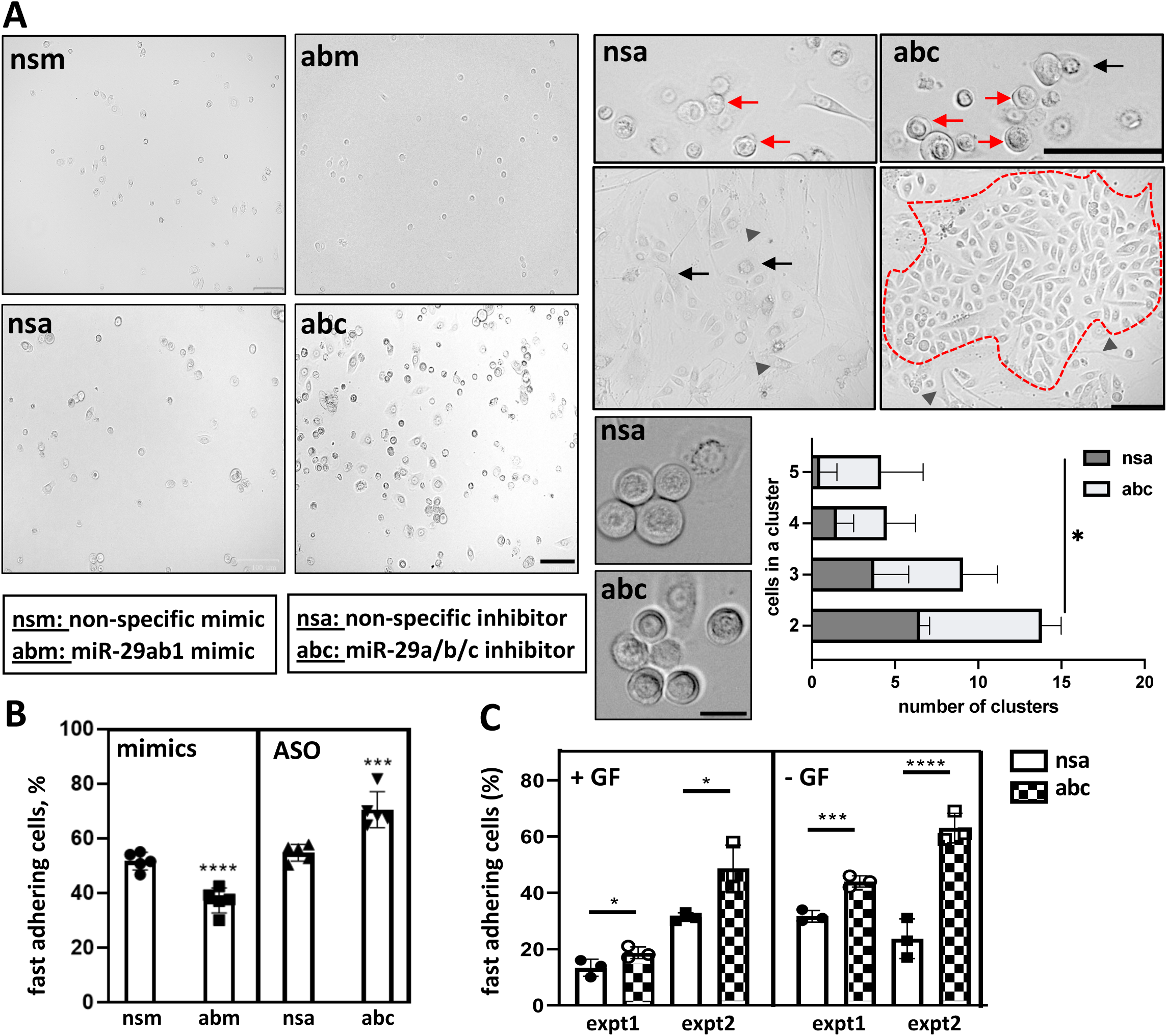
Loss of miRNA-29 enhances basal keratinocyte adhesion. Human primary keratinocytes were double transfected with 200nM of miR-29 inhibitors (abc) or non-specific antisense (nsa) or 50nM of miR-29 mimics (abm) or non-specific sense oligo (nsm) and the fast adhering cells were separated using fibronectin coated plates. **(A)** Representative images of the cells from each treatment. The arrows indicate clusters of more than two cells attached to the matrix, with red arrows indicating clusters with a potential for clonogenicity. Arrowheads show keratinocytes that acquired a spindle-like morphology with low nuclear-to-cytoplasm ratio indicating lost potential to form a holoclone. Scale bar 75µm (left four panels) and 50μm (four panels on the right). **(B)** Number of cells in clusters were counted in each treatment and plotted (n=4), paired t-test. Total cell number was determined using PrestoBlue, followed by one-way ANOVA. **(C)** Adhesion assay as done in (A) was performed for anti-miR-29 with +/- growth factors, and total cell numbers determined using PrestoBlue (n=3). *P < 0.05 or **P < 0.01 or \*\*\**P* < 0.001 or \*\*\*\**P* < 0.0001, two-way ANOVA followed by Šídák’s multiple comparison.

It was thus very important to identify the mechanisms released by miRNA-29 suppression in fast-attaching cells. To do this, we first sequenced the total RNA of fast and slow adhering keratinocytes along with respective controls, as explained in Suppl. Fig S4A. In an initial exploratory data analysis, we performed principal component analysis (PCA) of the raw data which showed the differences between the transcriptome of these cell populations (Suppl. Figure S4B). Indeed, the EnrichR analysis confirmed the aqusition of many more biological pathways (BP), cellular compartments (CC), and molecular functions (MF) in fast adhesring cells as a result of miRNA-29 removal (compare Suppl. Figure 5A and 5B). Protein binding in response to stimulus in the cytoplasm most significantly contributed to fast adhesion upon inhibition of miRNA-29 (Suppl. Figure 5C). It suggests that removal of miRNA-29 releases the expression of molecules that activate adhesion though cascades of signalling, starting from the transmembrane receptor proteins interacting with the second messengers and engaging more collateral mechanisms. We could not identify any previously known adhesion mechanism strongly enriched in fast miRNA-29 knock-down cells. This suggests that miRNA-29 represses basal adhesion through multiple nodes of regulation. Molecules in these nodes could be both direct and indirect targets of miRNA-29, forming a complex network. Thus, it is necessary to identify which mRNAs are bound and downregulated by miRNA-29 directly. Such ‘master regulators’ of adhesion are important because they may comprise a new network regulated in a concord rather than a single adhesion molecule or a pathway targeted by the miRNA.

### miRNA-29 directly targets new master regulators of adhesion in human skin

To identify the mRNAs bound by miRNA-29, we used miRNA-CLIP with the miRNA-29 probe, developed and tested in HEK293T cells and in immortalized human keratinocytes. We performed miRNA-29-CLIP in three types of primary cells isolated from human skin. The first type was the human follicular keratinocytes (HFK), the epidermal stem cells isolated from the sculp. The second type was the human interfollicular keratinocytes (IFK) and the third type was the human dermal fibroblasts (DF), all types isolated from healthy donors. Biotinylated miRNA-29 probe was transfected into HFKs, IFK, and DFs, crosslinked to mRNA targets, and the AGO2 pull down was perfomed to isolate a functional RNA-induced silencing complex (RISC). The RNA-miRNA duplexes were then extracted from the AGO2-RISC fraction and precipitated on streptavidin-coated beads, isolating the RNAs bound by the miRNA-29 probe. The resulting cDNA libraries thus contained the RNAs bound and targted to the RISC by miRNA-29. Before proceeding towards the sequencing of the libraries, qPCRs were performed to ensure the enrichement of known and predicted miRNA-29 targets in the probe fraction compared to the AGO2-RISC (IP1) and input. A significant enrichment of miRNA-29 targets was observed in all three cell types tested (Suppl. Figure S6A-C). Sequences of miRNA-29-CLIP libraries were analysed based on the number of reads and their enrichment. The predicted targets of miRNA-29 showed a significant enrichment in all three cell types (Suppl. Figure S6D). The HFK and IFK miRNA-29 targetomes shared more than 50% of their targets with each other meanwhile less than a half was shared between targetomes of keratinocytes and fibroblasts (Figure 4A). This result shows different miRNA-mRNA pairs even in the cells of the same or related origin and function (HFK and IFK). To determine the representation of miRNA-29 targets in all RNAs captured by the probe, we used two prediction tools: (1) TargetScan, which predicts targets based on evolutionary conservation (2) microT-CDS, which uses a machine-learning approach to detect miRNA-target interaction criteria from experimental data^22–24^. As expected, miRNA-29a-3p targets had the highest representation (∼60%, Figure 4B), correlating to the abundance of this miRNA-29 compared to other family members, and most of miRNA-CLIP mRNAs were predicted targets of miRNA-29 family for all three cell types (Figure 4C).

**Figure 4.**
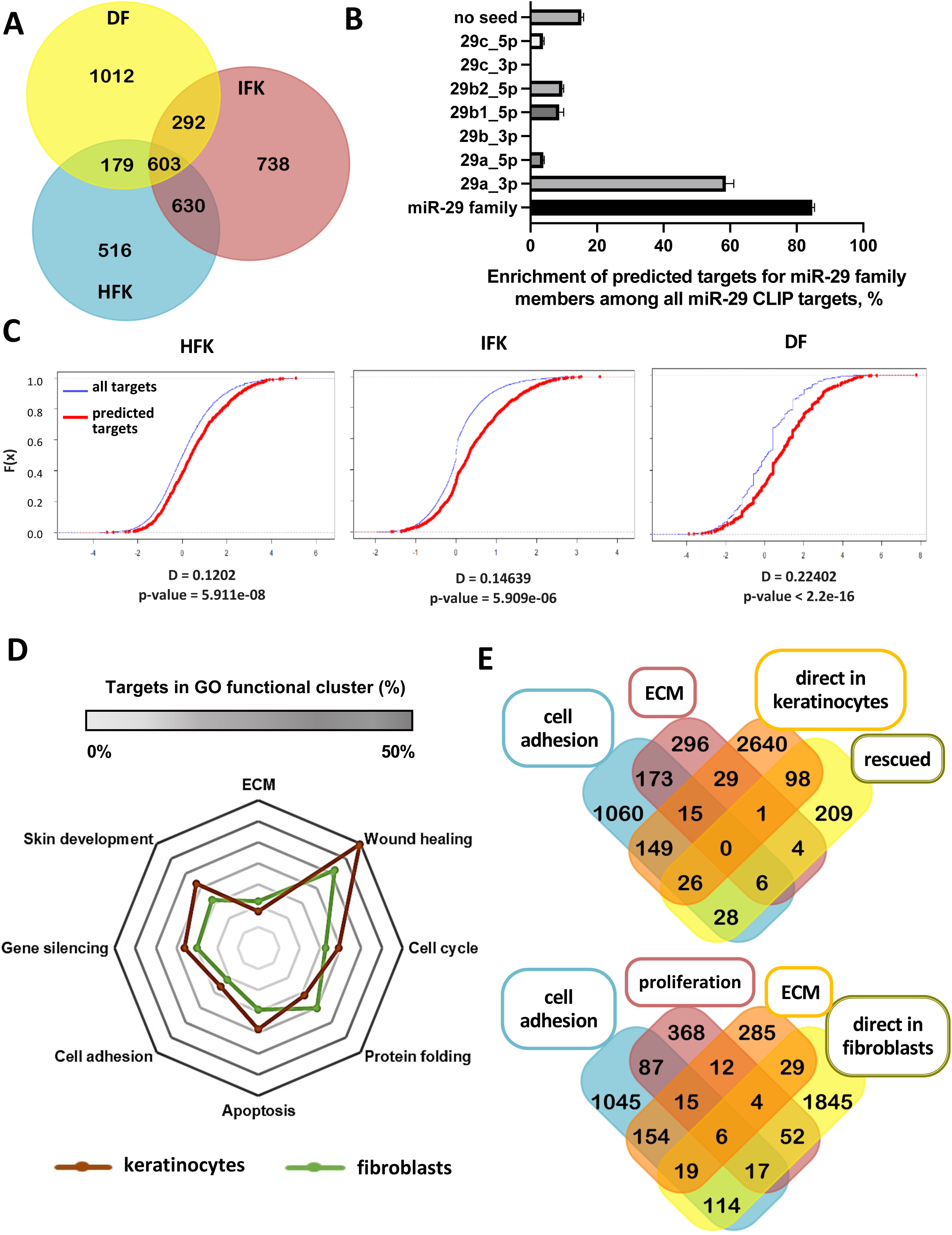
miRNA-CLIP successfully identifies miRNA-29 targetome in primary skin cells. **(A)** Venn diagram of overlap between the miR-29 targetomes (identified using miRNA-CLIP) of skin cells. **(B)** Percentage of all miRNA-29 family predicted targets captured by the miRNA-29 probe in all 3 miRNA-CLIPs. **(C)** Empirical Cumulative Distribution Function (ECDF) plotted the distribution of all miRNA-29-CLIP targets (blue) with predicted targets (red) against log2FC with distance and the p value for each of the cell types. This shows target enrichment achieved by the CLIP method vs. all target prediction. **(D)** Radar plot with the percentage representation of keratinocytes (HFK+IFK) and fibroblasts (DF) targetome in selected GO terms. **(E)** *Top:* Venn diagram showing the overlap of targets in GO term cluster for cell adhesion and ECM, with miRNA-29-CLIP targets (direct) and mRNAs rescued by miRNA-29 inhibition in keratinocytes. *Bottom:* the overlap of targets in GO term clusters for cell adhesion, proliferation, and ECM, with miR-29-CLIP targets in fibroblasts.

To compare miRNA-29 targetomes in HFK, IFK, and DF, we performed cluster analyses of miRNA-29 CLIP based on GO terms. When we chose the GO relevant to skin repair and barrier functions and accessed the percentage of miRNA-29 targets in them, we found that miRNA-29 targets cluster the most in the wound healing category for both keratinocytes and fibroblasts (Figure 4D). Comparing the fast adhesion targets rescued by miRNA-29 KD to direct targets of miRNA-29 from CLIP resulted in 26 miRNA-29-mRNAs with the function in adhesion (Figure 4E). Interetingly, zero mRNAs belonged to the ECM-related adhesion in keratinocytes, suggesting that miRNA-29 targeted mRNAs endogenous to keratinocytes, which were not involved the direct regulation of the ECM. The overlap of miRNA-CLIP targets from fibroblasts with GO clusters showed that in contrast to keratinocytes, miRNA-29-mediated adhesion in fibroblasts is connected to proliferation and deposition of the ECM (Figure 4E).

In line with the data, cell adhesion was the most significant category in keratinocytes and fibroblasts when all miRNA-29-CLIP targets were analysed by DAVID (Figure 5).

**Figure 5.**
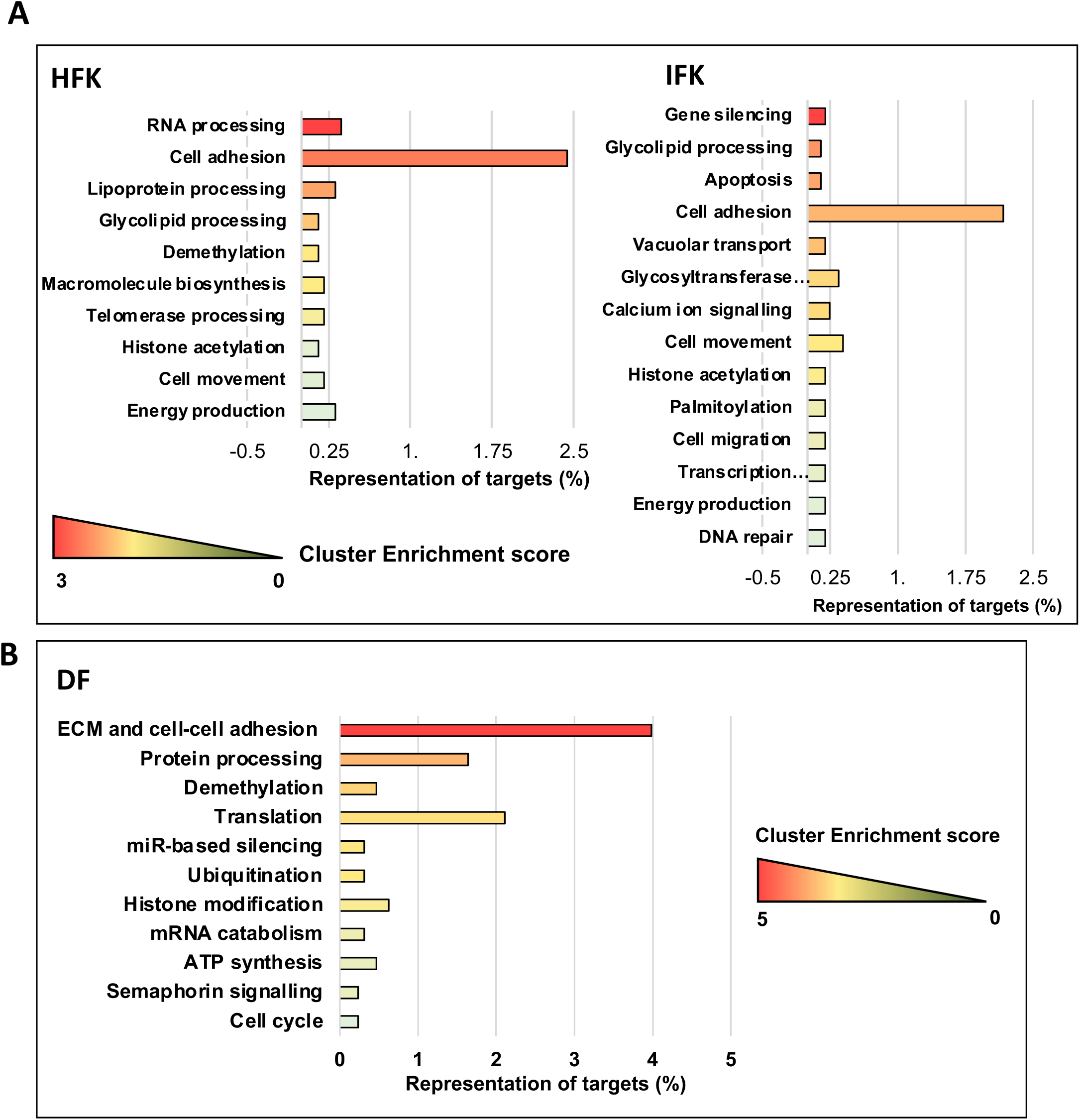
Functional analysis of miR-29-CLIP targetome in primary skin cells using DAVID. (A, B) Functional analysis of the miR-29-CLIP targetome using DAVID online tool for human follicular keratinocytes (HFK), interfollicular keratinocytes (IFK), and dermal fibroblasts (DF). Number of miR-29 targets represented in each of the cluster is on the x axis and the colour of the bars represent the cluster enrichment score.

EnrichR showed mRNA clusters describing skin-related functions and suggested that focal adhesion was more dependent on miRNA-29 in follicular keratinocytes compared to the interfollicular cells (Suppl. Fig. S7). miRNA-29 was also directly involved in regulating the interaction with immune cells, cell cycle, and lipid metabolism in keratinocytes and fibroblasts, with the IFK having the highest representation of lipid metabolism, and DF showing miRNA-29 function in the cell cycle and DNA methylation (Suppl. Fig. S7). Thus, many targets of miRNA-29 appeared in multiple pathways during GO annotations, confirming our hypothesis about miRNA-29 regulating cluster functions rather than individual adhesion pathways.

To further test this, we manually searched for the functions of miRNA-29 target mRNAs in each of the relevant pathways, choosing the mRNAs with a known role and measuring their expression following miRNA-29 inhibition. We confirmed *PLAUR, FERMT2, LYN*, and *TXNIP* as direct targets of miRNA-29 in the total and fast adhering populations of primary human keratinocytes (Figure 6A, B). In addition, *BCL2L11* and *NEDD9* were specifically rescued by miRNA-29 inhibition in fast adhering keratinocytes (Figure 6B).

**Figure 6.**
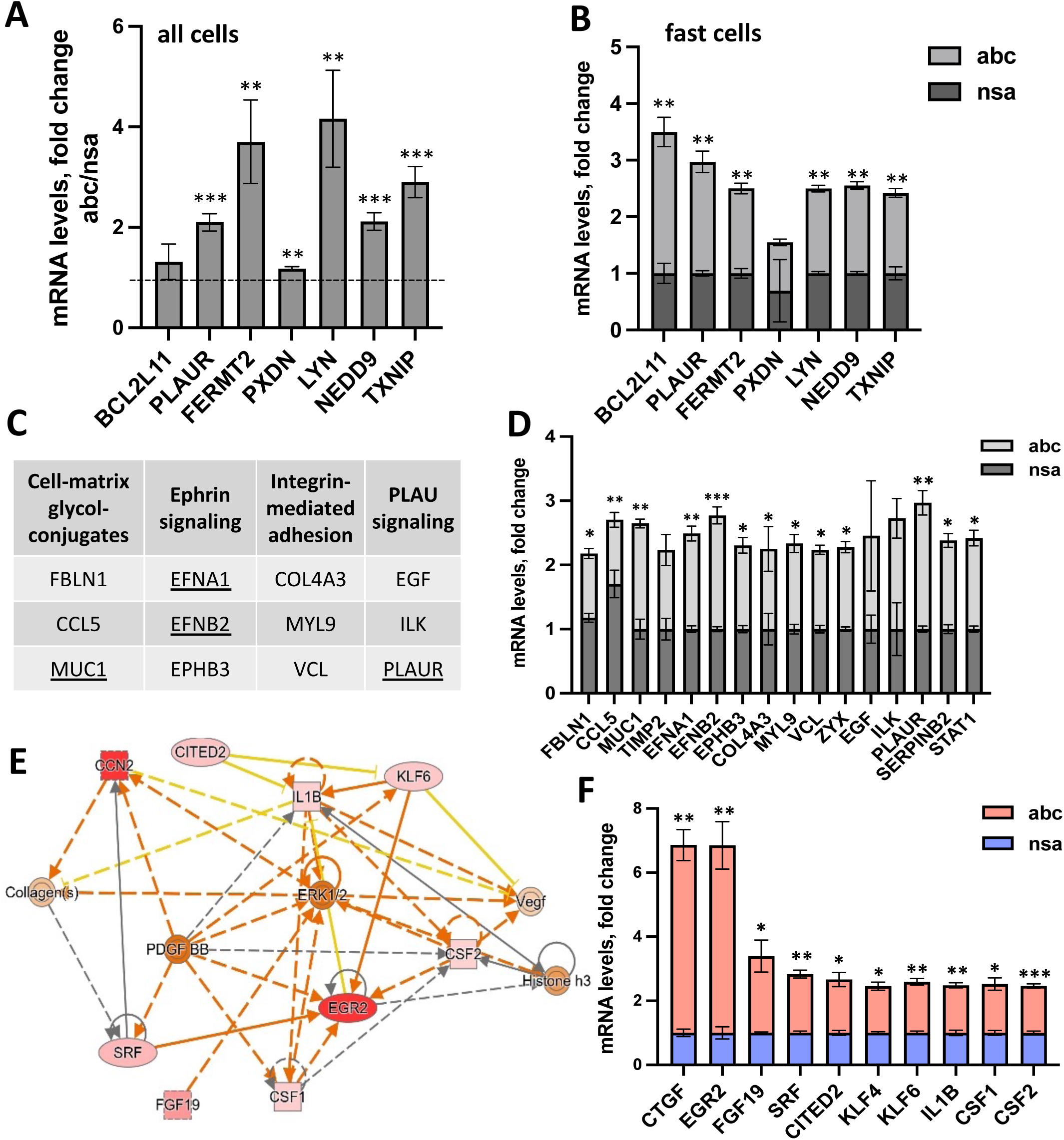
Identifying factors that contribute to enhanced adhesion in miR-29 knock-down cells. Fold change of selected known growth factors and regulators of adhesion in all keratinocytes **(A)** or fast adhering keratinocytes **(B)** transfected with miRNA-29 inhibitors (abc) versus control (nsa) and quantified using qPCR. Baseline expression of the nsa-treated samples is set to 1 in (A) and a two-way ANOVA followed by Šídák’s multiple comparison test is performed resulting in P value * < 0.05, **< 0.01, *** < 0.001. mRNAs that represent 4 pathways related to epithelial cell adhesion **(C)** were selected and changes in expression after miRNA-29 inhibition were plotted in **(D).** The most significantly rescued mRNAs are underlined in **(C). (E)** Ingenuity Pathway Analysis (IPA) generated heat map of diseases and functions from the fast keratinocytes (abc) versus control (nsa). **(F)** Regulators of TGF-ß/ß-catenin pathway were selected and their fold change in fast abc versus control nsa were plotted (N=3), unpaired t-test: *P < 0.05, **P < 0.01, ***P < 0.001.

Further analysis of the daughter categories of cell adhesion pathways (Suppl. Table 1) identified miRNA-29-dependent regulators of early cell adhesion that could be grouped into four pathways that function in basal keratinocytes (Suppl. Table 2 and Figure 6C). *MUC1, EFNA1, EFNB2* were most significantly upregulated in the fast cells after inhibition of miRNA-29 (Figure 6D), suggesting the involvement of cell-matrix glycol-conjugates and ephrin signaling in fast adhesion. In addition, we analyzed miRNA-29-CLIP targets in TGF-β and β-catenin pathways (Figure 6E). IPA identified more regulators of miRNA-29-controlled adhesion, namely, *CTGF, EGR2, SRF, KLF6, IL1B*, and *CSF2* (Figure 6E, F). These results confirmed that miRNA-29 controlled adhesion through the pathways with distinct functions^25^. When upregulated, these pathways could enhance basal adhesion together, each contributing only a part of the miRNA-29-controlled mechanism. To our knowledge, this is the first time when the regulators of adhesion are identified as directly regulated by a miRNA as a ‘master’ complex between human epidermal and dermal compartments, a cross talk which requires further investigation due to its high importantce in functionality and repair of human skin.

### miRNA-29 regulates adhesion through ECM deposition and paracrine mechanisms

Epidermal keratinocytes produce their own matrix as they attach on uncoated surfaces, whereas *in vivo* this process is coordinated with the underlying fibroblasts of the dermis. To further elucidate the dermal contribution to the miRNA-29-mediated regulation of adhesion, we used fluorescently labeled anti-sense oligos (ASOs) to inhibit miRNA-29 in fibroblasts. Based on previously reported functions of miRNA-29 in suppressing fibrosis, we expected to see an enhanced differentiation of fibroblasts to myofibroblasts. To our surprise, loss of miRNA-29 did not lead to appearance of myofibroblasts in culture and even suppressed fibroblast to myofibroblast conversion in response to TGFβ1 (Figure 7A). Lower levels of smooth muscle actin (aSMA) in miRNA-29 knock-down fibroblasts were confirmed by western blot (Figure 7B). We also observed a decrease in fibronectin secretion by fibroblasts treated with miRNA-29 ASO (Figure 7B). At the same time, inhibition of miRNA-29 significantly increased the number of fibroblasts at day 3-5 following transfection with ASO (Figure 7C). It is worth noting that the miRNA-29 ASO localized to rounded (as opposed to differentiated, aSMA-positive) fibroblasts, suggesting that inhibition of miRNA-29 supports their de-differentiated state, allowing proliferation (Figure 7A, C) and possibly, resulting in higher deposition of the ECM due an increase in cell number.

**Figure 7.**
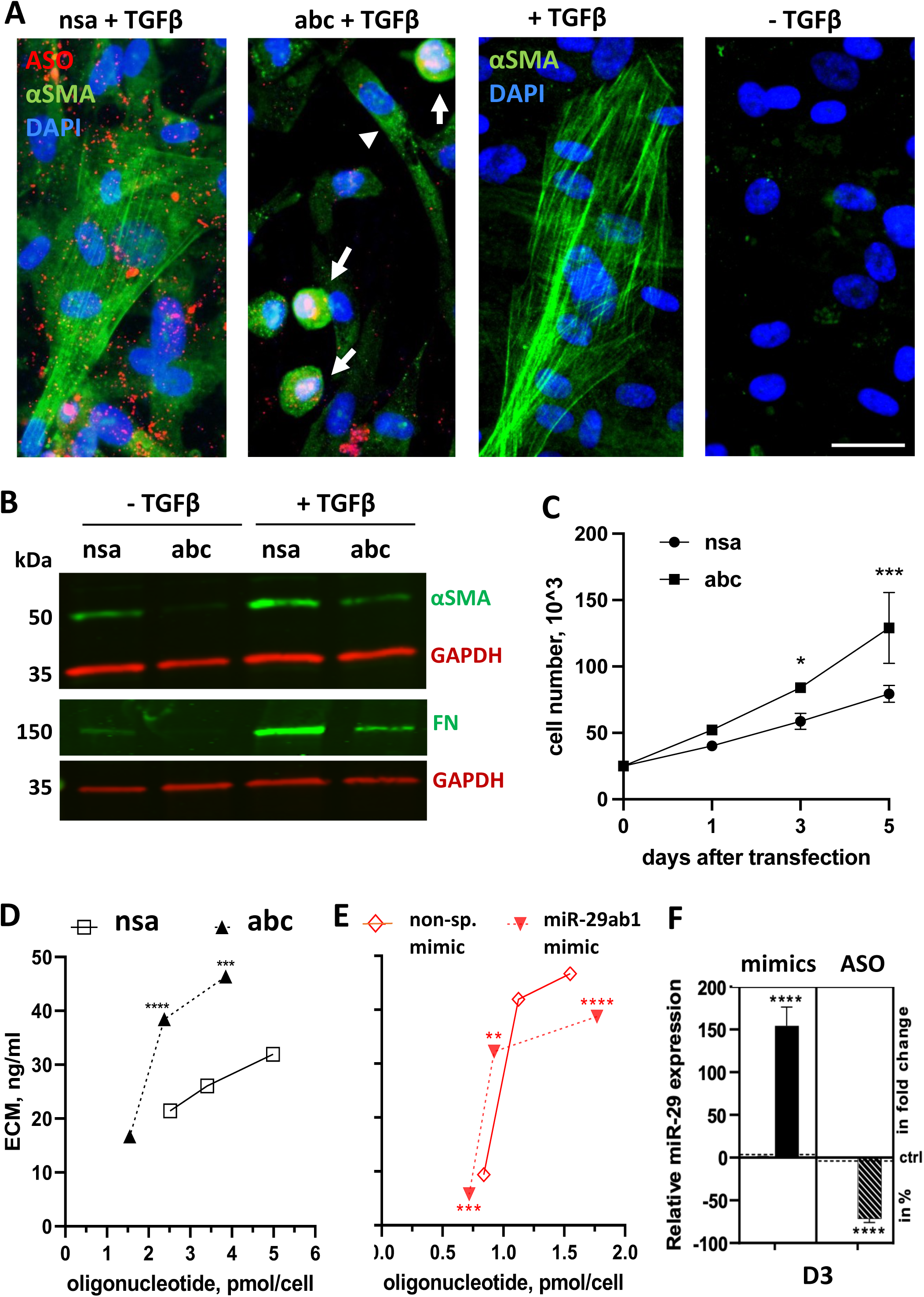
miRNA-29 regulates proliferation and extracellular matrix deposition in dermal fibroblasts. **(A)** Expression of αSMA was assessed by immunofluorescence in human primary DF transfected with anti-sense miRNA-29 (abc) or non-specific (nsa) and treated with 1ng/ml TGF-β1. Representative images are shown for nuclei staining with ASO in red, DAPI in blue, and αSMA in green. Arrows indicate localization of ASO inside the round-shaped fibroblasts with miRNA-29 LOF. Arrowheads point to a spindle-shaped cell not transfected with ASO. Scale bar is 25μm. **(B)** Expression of αSMA and fibronectin was carried out by western blot in transfected fibroblasts after the treatment with TGF-β1. GAPDH was used as a loading control. **(C)** Growth rate of fibroblasts was determined following miRNA-29 knockdown compared to non-specific oligo. **(D and E)** DF were double transfected anti-sense miR-29 (abc) or non-specific (nsa) or miR-29ab mimic (abm) or non-specific mimics (nsm) and the extracellular matrix (ECM) deposited by the cells was extracted by pepsin and quantified using BCA at 1,3, and 5 days post transfection. **(F)** Relative levels of miR-29 was measured after the second transfection to assess the knock-down/up levels. 2-way ANOVA followed by Šídák’s multiple comparison test. *P < 0.05 or **P < 0.01 or \*\*\**P* < 0.001 or \*\*\*\**P* < 0.0001.

Indeed, loss of miRNA-29 resulted in significantly more ECM secreted by the fibroblasts (Figure 7D) whereas addition miRNA-29 suppressed ECM (Figure 7E). While this result is overall consistent with the reported data on the miRNA-29-mediated fibrotic phenotype in skin^16^, lung^26^, kidney^27^, and liver^28^, it suggests that the mechanism of miRNA-29-regulated ECM deposition is not merely due to the de-repression of ‘classic’ pro-fibrotic targets. Rather, it may be the result of higher proliferative capacity of fibroblasts with low levels of miRNA-29. Gain of miRNA-29 function, however, may directly suppress the pro-fibrotic targets, such as collagens I and III, resulting in less ECM (Figure 7E, F).

To understand the mechanism by which inhibition of miRNA-29 induces proliferation and ECM deposition in dermal fibroblasts, we went back to the miRNA-CLIP targets involved in ECM deposition, proliferation, and adhesion (Figure 4E and 5B). Compared to Venn diagram for keratinocytes, fibroblasts had more direct targets of miRNA-29 represented in the ECM deposition category (one *vs.* 10, correspondingly, Figure 4E). We chose mRNAs that overlapped between ECM deposition and cell proliferation and were direct miRNA-29 targets from the DF miRNA-CLIP (Figure 4E). We compared their expression in fibroblasts with suppressed miRNA-29 and the control, confirming *SPARC*, *LOXL2*, *FERMT2*, *PXDN, COL4A1,* and *COL4A2* as direct targets of miRNA-29 (Figure 8A). Unlike in keratinocytes, *BCL2L11* expression remained unchanged and *PLAUR* was significantly downregulated (Figure 8A). We confirmed the regulation at the protein level for the highest upregulated mRNAs of SPARC, FERMT2, and COL4A1/2 (>1.5 fold increase in the miRNA-29 ASO treated fibroblasts, (Figure 8A, B). Inhibition of miRNA-29 significantly increased the expression of SPARC protein as early as 5 hours post transfection and remained elevated for 72 hours whereas FERMT2 protein responded to miRNA-29 inhibition only after 24 hours, probably reflecting differences in mRNA and protein turn-around time (Figure 8B-D). Consistently, fibroblasts expressing higher levels of FERMT2 had a more rounded morphology compared to the star-shaped control cells (Figure 8C).

**Figure 8.**
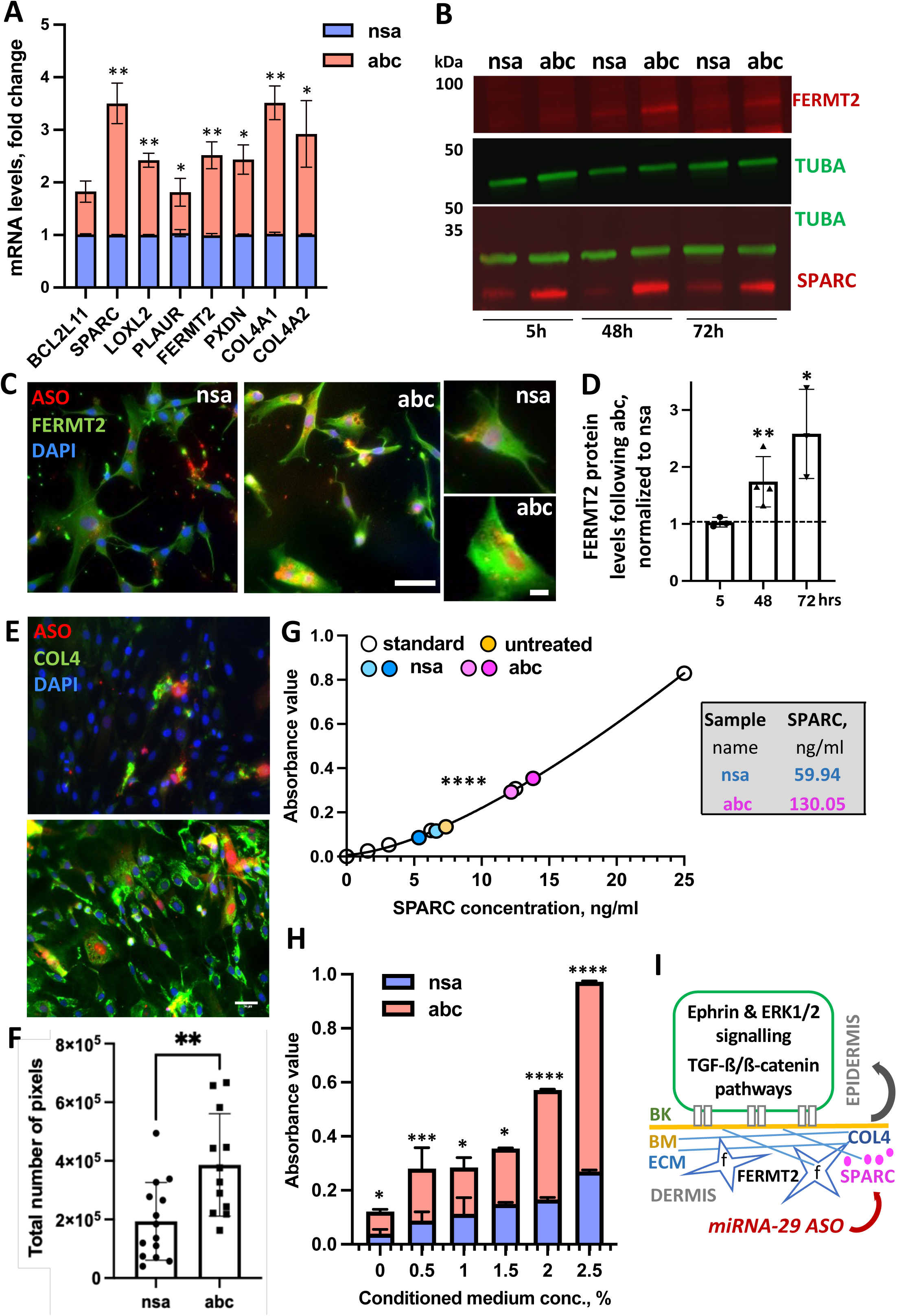
miR-29 regulates ECM, proliferation, and adhesion through SPARC, FERMT2, and COL4. **(A)** Selected miR-29 target genes represented in ECM and cell-cell adhesion cluster were confirmed by qPCR in human primary dermal fibroblasts. **(B)** Expression of SPARC and FERMT2 was assessed by western blot with tubulin as a loading control. **(C)** Representative images for FERMT2 (green), anti-miR-29 oligo (red) and nuclei (DAPI). Scale bar = 25μm. **(D)** Quantification of FERMT2 expression from western blots at 5 hr, 48hr and 72 hr post transfection, N=3, unpaired t-test. *P < 0.05 or **P < 0.01. **(E and F)** Representative images **(E)** and quantification of collagen IV **(F)** using ten images per condition. Unpaired t-test, **P < 0.01. Scale bar = 35μm. **(G)** Soluble SPARC was quantified by ELISA in the medium conditioned by miRNA-29 ASO (abc) and control (nsa) ASO. Graph shows direct ELISA results where medium samples were diluted for precise quantification. The actual levels of SPARC are shown as numbers next to the graph. Both one-way and two-way ANOWA followed by Šídák’s multiple comparison tests were performed, showing highly significant increase in secreted SPARC, ****P<0.0001. **(H)** Adhesion of keratinocytes was quantified on the increasing concentrations of conditioned media from miRNA-29 ASO (abc) or control fibroblasts (nsa). N=3, unpaired t-test, *P < 0.05 or **P < 0.01. Please note the absorbance value coming from the PrestoBlue reagent reduced by alive cells and thus, corresponding to their number. **(I)** Schematics of the interaction between proposed players of enhanced keratinocyte adhesion regulated through miRNA-29. BK – basal keratinocyte, BM - basal membrane, ECM – extracellular matrix, f – fibroblasts, and ASO - antisense oligonucleotides.

Our results showed the expression of *COL4A1* and *COLA4A2* increased by than 2- and 1.75-fold, respectively, confirming previous findings that these two mRNAs are direct targets of miRNA-29 ^29^. The increase in pan-COL4 protein observed in the culture staining was quite striking (Figure 8E, F), and confirmed regulation of collagen IV by miRNA-29 in fibroblasts.

SPARC (secreted protein acidic and rich in cysteine) is an extracellular glycoprotein expressed in elevated levels in actively proliferating cells, with indespensible function during wound healing^30^. SPARC has many post-tranlational modifications resulting in various isoforms, with the shortest one (32 kDa, Figure 8B) being secreted and involved in cell-ECM interactions^31^. To validate the localization of increased levels of SPARC observed in fibroblasts with inhibited miRNA-29 (Figure 8B), we performed ELISA assays on the medium conditioned by miRNA-29 ASO-treated DFs vs. control ASO. As expected, SPARC was secreted by fibroblasts ^32^, and inhibition of miRNA-29 significantly increased the concentration of SPARC in the medium compared to control (Figure 8G).

Finally, we tested if these changes in the fibroblast secretome induced by miRNA-29 inhibition can regulate keratinocytes adhesion via paracrine mechanisms. We collected the medium produced by DFs, which were transfected with miRNA-29 ASO or control oligos. The conditioned medium was concentrated by lyophilization and added at increasing concentrations to untransfected keratinocytes as a substrate for the adhesion assay. We found that the number of rapidly adhering keratinocytes increased in the medium produced by miRNA-29 ASO-treated DFs compared to the control ASO conditioned medium (Figure 8H). This result demonstrates that the inhibition of miRNA-29 can also induce adhesion of basal keratinocytes through the paracrine signaling from the underlying dermal fibroblasts (Figure 8I).

Important to the mechanism of miRNA-29-mediated regulation, *FERMT2*, *COL4A1*, *COL4A2, SPARC, and LOXL2* (Figure 8) showed high miTG score indicating strong binding of the target 3’UTRs to the miRNA-29 ‘seed’ (Suppl. Figure S8). Thus, we report direct targets of miRNA-29 that are endogenous to dermal fibroblasts and regulate proliferation, ECM depostion, and adhesion through the key sigaling and paracrine mechanisms. We demonstrate that miRNA-29 regulates adhesion in the skin by direct binding and regulation of mRNAs encoding proteins from many signaling pathways, including ephrin, MAPK/ERK, and TGFβ/β-catenin. We show that inhibition of miRNA-29 increases basal adhesion of keratinocytes, supporting growth of the human epidermis *ex vivo*, and correlating to low levels of miRNA-29 in basal epidermis in mouse wounds *in vivo.* These findings identify miRNA-29 as a target for ASO-mediated therapy, which has a potential to increase regenerative capacity of epidermis and dermis during skin repair.

## DISCUSSION

### miR-CLIP indentifies direct, cell-specifcic, and functional mRNA targets of miRNA-29

In this study, we aimed to investigate new functions of miRNA-29 in the skin during its normal growth and regeneration. We chose the human skin as a model to unveil cell-specific miRNA-29 targetome in primary keratinocytes and fibroblasts, two major cell types of the skin, during their growth and differentiation in 2D and 3D models ex vivo. A common approach identifies expression levels of target mRNAs after partial removal or an overexpression of a miRNA, followed by a target prediction through the ‘seed’ sequence matching of the 3’UTR. Similarly, a pull-down of the miRNA-29 mimic labeled with biotin alone results in a capture of complementary RNA sequences outside of AGO2-RISC, whereas the RNA immunoprecipitation (RIP) crosslinking precipitation (CLIP) of Ago2-RISC does not reveal which miRNAs interacted with each of the isolated binding sites. Our approach, however, used low concentrations of the miRNA stem-loop precursor, covalently linked to biotin and psoralen in nucleotide positions, which did not change the processing of the stem-loop into the functional miRNA but allowed the tandem pull-down of AGO2-biotin complexes of miRNA-29 and mRNAs, crosslinked near the ‘seed’ sequence^1^.

### miRNA-29 inhibition confers fast adhesion through multiple pathways

Cell adhesion plays an important role in skin regeneration and wound healing. Upon excisional wounding, the keratinocytes migrate over the fibronectin (FN) in the wound bed matrices for wound closure^33^. Also, FN reorganization in basal membrane influences focal adhesion which is essential in cell attachment and survial^34^. Keratinocytes secret basement membrane components in response to an injury^35^ and it is interesting to note that basal membrane components like nidogen-2, netrin-1, netrin G1, and agrin are expressed more in the fast adhering keratinocytes treated with miRNA-29 ASO (Suppl. File 1).

Adhesion molecules are proteins on the cell surface that are involved in the interactions between cell surface and the ECM. Adhesion molecules include cadherins (subgroups E, N, P, M), integrins, selectins, and the immunoglobulin gene family. We detected increased expression of less studied cell adhesion molecules such as smoothlin, NCAM, MCAM, EPCAM, MADCAM1, CADM4, Nectin 2, Nectin 4, CEACAM and L1CAM, ICAM-1, ICAM -5 and BCAM, on the fast population under miRNA-29 control (Suppl. Table 3). All these molecules have been associated with increased cell adhesion and proliferation^36, 37^. A regulator of keratinocyte differentiation predominately expressed in the basal layer, TXNIP^38^, was also upregulated in fast adhering population of miRNA-29 KD cells.

Indirect suppression of proteins in response to miRNA-29 inhibtion may also help the cells to adhere faster. For example, α-actinins, highly related members of the spectrin superfamily, are ubiquitously expressed in most cells of the body. Depletion of the focal adhesion protein α-actinin enhances force generation in initial adhesions on fibronectin, but impairs mechanotransduction in a subsequent step, preventing adhesion maturation^39^. It is interesting to note that actinin 2 and 4 are down-regulated in fast adhering cells after miRNA-29 ASO. Finally, the absence of growth factors increased the proportion of rapid adhering cells after miRNA-29 inhibition, suggesting the possibility of a growth factor-independent mechanism driving the miRNA-29-regulated adherence.

### The regulation of extracellular matrix by miRNA-29 is more complex and reversible than previously thought

Increased ECM deposition by fibroblasts is often associated with myofibroblasts activation and TFGβ1 signaling. Many studies have reported the effect of TGFβ1 on the endogenous expression of miRNA-29, but the effect of miRNA-29 inhibition on TGFβ1 in primary dermal fibroblasts had not been reported^40^. Initial ECM assembly of latent TGF-β1 binding protein is strongly dependent on the presence of FN ^33, 41^. Our results indicate that the increase in ECM synthesis in miRNA-29 ASO-treated cells goes through a myofibroblast-independent mechanism. These observations indicate that the miRNA-29 family can regulate both TGF-β-dependent and -independent pathways.

From the direct targets of miRNA-29 from the DF miRNA-29-CLIP, four mRNAs involved in ECM deposition, cell adhesion, and proliferation specifically stood out. Kindlin-2/FERMT2, SPARC, PXDN and LOXL2 are released into the extracellular space and are associated to collagen fibril organization, contributing to ECM homeostasis^42^. FERMT2 activates integrins leading to FN binding and adhesion, and subsequently assembles an essential signaling node at newly formed adhesion sites in a talin-independent manner^39^. Its major role is to regulate the activation of the integrins and is essential for cell shape.

Peroxidasin PXDN catalyzes the two-electron oxidation of bromide by hydrogen peroxide and generates hypobromite as a reactive intermediate in the formation of cross-links between methionine and hydroxylysine residues of collagen IV hexamer. This directly contributes to the collagen IV network-dependent fibronectin/FN and laminin assembly, which is required for full extracellular matrix (ECM)-mediated signaling. SPARC binds a subset of ECM proteins, regulating ECM assembly through collagen interaction with the cell surface, and can be suppressed by miRNA-29^42, 43^. Lysyl oxidases (LOX) is the family of extracellular fibrillar collagen crosslinking enzymes that mediate the progression from fibrils to fibers and it is integral in maintaining ECM organization. LOXL2 has been reported to be regulated by miRNA-29 during cancer progression^44^. Our results identified and confirmed previously reported endougenous tragets of miRNA-29 in fibroblasts and suggested that SPARC, FERMT2, PXDN, and LOXL2 act in a concerted fashion to facilitate *the novo* collagen incorporation into fibrils and the ECM assembly.

Taken together, our results revealed the direct *in vivo* targetome and functions of miRNA-29 in three types of cells isolated from human skin. We uncovered new mechanisms of miRNA-29-regulated cell adhesion and deposition of the ECM, and showed that these can be significantly enhanced by the inhibiton of miRNA-29. Our results demonstrate that changes in a single miRNA can influence different pathways in a cell-type specific manner. When studied in the context of the organ, however, these pathways remarkably converge to regulate the most essential functions, such as restoration or the barrier formation in the skin. Since miRNAs can be up- or -downregulated by short oligonucleotides, our findings open exciting opportunity to regulate these functions by new therapeutic interventions.

## MATERIALS AND METHODS

### Mouse maintenance and experimentation

Mice were maintained under specific pathogen-free conditions and obtained food and water *ad libitum*. Mouse maintenance and experiments with mice were approved by the UK veterinary authorities. Two 6mm wounds were introduced into the dorsum of each mouse using a sterile biopsy punch and wounds were collected using an 8mm biopsy punch post-mortem.

### Cell culture and transfection of oligonuclotides

Primary human cells were isolated from neonatal foreskin representing interfollicular keratinocytes (IFK) and dermal fibroblasts (DF) obtained from PeloBiotech and maintained in serum-free keratinocyte growth media (PeloBiotech) at 37 °C, 5% CO2 on Coating Matrix Kit Protein (Gibco) coated flasks. Human follicular keratinocytes (HFK) were isolated from the hair of healthy donors and maintained as IFKs. The cells were used at passage 3-4 in all experiments. DF were maintained in fibroblasts growth medium kit (PeloBiotech) at 37 °C, 5% CO2. The miRNA-29-CLIP probe, 409 was diluted in Optimem (Life Technologies, 7.5 nM (keratinocytes) or 5 nM (fibroblasts) final concentration and transfected with RNAiMax (Life Technologies) to 15-cm dishes of cells at 90% confluency and incubated for 24 h. A mock transfection was set up with the same condition as above without the probe. Due to different endogenous levels of miRNA-29 in HFK, IFK, and DF (higher miRNA-29a/b1 expression detected in keratinocytes than in fibroblasts in our initial experiments), we used low concentrations of the probe, which would not elicit a strong gain-of-function response, 7.5nM for keratinocytes (IFK and HFK) and 5nM for fibroblasts (DF).

For stem-loop miRNA mimics and single-stranded antisense oligonucleotide (ASO) transfections, the cells were seeded at 70–80% and 50-60 % confluence, respectively (day 0) and incubated overnight at 37^0^C, 5% CO2. On day 1, the cells were transfected with miRNAVana miRNANA mimics or inhibitors (ASO) at a final concentration of 50nm and 200nM, respectively, using Lipofectamine RNAiMAX (Thermo Fisher Scientific) according to the manufacturer’s protocol. For the efficient inhibition of miRNA-29, the cells were re-transfected on day 3 and collected for the RNA analysis or functional assays on day 5. All transfections were performed in biological and technical triplicates. A complete list of used miRNA mimics, anti-miRNAs, cloning and Taqman qRT-PCR primers are provided in Suppl. Table 4.

### miRNA-29-CLIP, followed by qPCR, QC, and NGS library preparation

100 μl of Prot G Dynabeads (Life Technologies) per 15-cm dish was washed thrice with 1 ml of citrate-phosphate buffer (25 mM citric acid, 66 mM Na2HPO4, pH 5.0). The Ago2 antibody (50 μg per 100 μl of Prot G Dynabeads) was immobilized in a total volume of 500 μl of citrate-phosphate buffer by gentle rolling for 1 h at 4 °C. The beads were washed thrice with 1 ml of NP40 lysis buffer and blocked with 1 ml of NP40 lysis buffer containing BSA (10 μg/ml) for 0.5 h at 4 °C. After three washes with 1 ml of NP40 lysis buffer, the beads were resuspended in 50 μl of the lysis buffer.

20 µl magnetic streptavidin beads per 15 cm dish were prepared as per manufacturer’s protocol. The washed beads were blocked in binding and washing buffer with salmon sperm DNA (100 μg/ml), BSA (100 µg/mL) and heparin (100ng/mL) for 1 h at 4 °C on a rotator. The beads were washed five times with binding and washing buffer and resuspended in 50 μl of binding & washing buffer for pre-equilibration.

After probe transfection, cells were washed once with 10 ml of PBS, placed on ice and irradiated twice at 254 nm (CL-1000 UltraviolettCrosslinker, UVP) with 100 mJ, followed by irradiation at 365 nm with 150 mJ. After irradiation, cells were scraped in 1.5 ml ice-cold PBS, pelleted at 200g at 4 °C for 5 min and lysed in 1 ml of NP40 lysis buffer for 15 min on ice. The lysate was cleared at 14,000g at 4 °C for 15 min and incubated with the Ago2 (clone 9A11; Ascenion) antibody-coupled Prot G beads for 1 h at 4 °C with gentle rolling. The beads were washed five times with each 1 ml of IP wash buffer (50 mM HEPES, pH 7.5, 300 mM KCl, 0.05% NP40, 0.5 mM DTT, complete protease inhibitor (Roche)). After the last washing step, 200 μl digest buffer (100 mM Tris-HCl, pH 7.5, 150 mM NaCl, 12.5 mM EDTA) containing 100 μl proteinase K (recombinant PCR grade solution, Roche) was added to each sample and digested at 65 °C for 15 min. RNA from the proteinase K digest was isolated by chloroform/phenol/isoamyl alcohol (Life Technologies) with 20 μg/ml Glycoblue (Life Technologies). After centrifugation for 10 minutes at 12 000g, the RNA was precipitated from the aqueous phase using 600 ml of 100% ethanol and 20 uL 3M NaOAc at -20°C overnight. Then, the samples were centrifuged at 12,000 g for 30 minutes at 4 °C to pellet the RNA and was dissolved in 20 µl water.

For streptavidin affinity purification, Ago-pulldown samples were incubated with streptavidin beads for 30 min at 4 °C with gentle agitation. The beads were washed four times with binding and washing buffer and treated with DNAse (PCR grade, Roche) for 10 minutes at 37 °C. Beads were washed once with 1 ml of high-salt buffer. The beads were shaken twice with 50 μl of a solution of 95% formamide/10 mM aqueous EDTA, pH 8.2, for 2 min at 65 °C. The solutions were combined, filled with 100 μl of water and 20 μl of 3 N aqueous sodium acetate and purified by chloroform/phenol (Life Technologies) extractions. The aqueous phase was isolated, and RNA was precipitated with 1 ml of 100% EtOH at −20 °C overnight with the addition of 20 μg/ml Glycoblue (Life Technologies). Then, the RNA pellet was washed with 70% EtOH, centrifuged at 14,000g for 10 min at 4 °C and dissolved in 20 μl of water.

Library was prepared using Illumina stranded Total RNA prep ligation with Ribo-Zero plus following the manufacturer’s protocol. Input RNA, RNA isolated from Ago2 IP (miRNA-29-CLIP probe, 409 or mock) (ribo-depleted) and miRNA-CLIP–purified RNA (not ribo-depleted) were submitted for Illumina sequencing. A small proportion of miRNA-CLIP– purified RNA was analyzed by q-PCR to control for successful enrichment of a set of validated miRNA-29 targets the following way. After reverse transcription (Multi-Scribe Reverse Transcriptase Kit, Life Technologies) of the isolated RNA, enrichment of target genes as normalized to GAPDH was measured by qRT-PCR (TaqMan Fast Advanced Master Mix, ThermoFisher) on a QuantStudio 12K Flex Real-Time PCR system (ThermoFisher) following the manufacturer’s protocol. Quality and integrity of the RNA samples were assessed using a 4200 TapeStation (Agilent Technologies) and then libraries generated using the Illumina® Stranded mRNA Prep. Ligation kit (Illumina, Inc.) according to the manufacturer’s protocol. Briefly, total RNA (typically 0.025-1ug) was used as input material from which polyadenylated mRNA was purified using poly-T, oligo-attached, magnetic beads. Next, the mRNA was fragmented under elevated temperature and then reverse transcribed into first strand cDNA using random hexamer primers and in the presence of Actinomycin D (thus improving strand specificity whilst mitigating spurious DNA-dependent synthesis). Following removal of the template RNA, second strand cDNA was then synthesized to yield blunt-ended, double-stranded cDNA fragments. Strand specificity was maintained by the incorporation of deoxyuridine triphosphate (dUTP) in place of dTTP to quench the second strand during subsequent amplification. Following a single adenine (A) base addition, adapters with a corresponding, complementary thymine (T) overhang were ligated to the cDNA fragments. Pre-index anchors were then ligated to the ends of the double-stranded cDNA fragments to prepare them for dual indexing. A subsequent PCR amplification step was then used to add the index adapter sequences to create the final cDNA library. The adapter indices enabled the multiplexing of the libraries, which were pooled prior to cluster generation using a cBot instrument. The loaded flow-cell was then paired-end sequenced (76 + 76 cycles, plus indices) on an Illumina HiSeq4000 instrument. Finally, the output data was demultiplexed and BCL-to-Fastq conversion performed using Illumina’s bcl2fastq software, version 2.20.0.422 Unmapped paired-end sequences from an Illumina MiniSeq sequencer were output in BCL format and converted to FASTQ format using bcl2fastq v2.20.0.422, during which adapter sequences are removed. Resulting reads were between 34 and 75 nucleotides in length. Unmapped paired-reads of 74bp were interrogated using a quality control pipeline consisting of FastQC v0.11.3 (http://www.bioinformatics.babraham.ac.uk/projects/fastqc/) and FastQ Screen v0.13.0 (http://www.bioinformatics.babraham.ac.uk/projects/fastq_screen/). The reads were trimmed to remove any adapter or poor quality sequence using Trimmomatic v0.36 (PMID: 24695404); reads were truncated at a sliding 4bp window, starting 5’, with a mean quality <Q20, and removed if the final length was less than 36bp. The filtered paired-reads were mapped to the human reference sequence analysis set (hg38/GRCh38) from the UCSC browser, using STAR v2.5.3a. The genome index was created using the comprehensive Gencode v25 gene annotation (http://www.gencodegenes.org/). Normalisation and differential expression analysis was performed using DESeq2 v1.18.1 on R v3.4.0. Normalisation of paired-read counts and calculation of regularized log2 fold change, without estimation of gene expression dispersion was performed using DESeq2 v1.18.1 on R v3.4.0. Genes were annotated using information extracted using Biomart for Ensembl v85, which corresponds to Gencode v25 (http://jul2016.archive.ensembl.org/). Unspliced sequence (UTR, exon, introns) was extracted using Biomart for Ensembl v85 (http://jul2016.archive.ensembl.org/). The resulting FASTA file was scanned using a Perl script to identify the regular expression matches to seed motifs on forward and reverse strands.

### Functional analysis of NGS results

The miRNA-29 targets captured by the probe were divided into strong, weak, and no seed binding based on their miRNAANDA scores and classified based on the target expression levels in HEK293T cells. The DAVID platform v6.8 functional annotation clustering tool was employed with UP_KEYWORDS, UP_SEQ_FEATURE, COG_ONTOLOGY, BBID, BIOCARTA, KEGG_PATHWAY, INTERPRO, PIR_SUPERFAMILY, SMART, GOTERM_BP_DIRECT, GOTERM_CC_DIRECT and GOTERM_MF_DIRECT as clustering features. Clusters containing enriched terms were sorted by size and given the nomenclature based upon the terms included. Predicted target transcripts of miRNA-29 were obtained from micro-TDS tool in DIANA website. The predicted miRNA-29 target transcripts with a z-score of above 0.3 were used for further comparison.

Ingenuity Pathway Analysis (IPA) was used to do comparative analysis of fast adhering and slow adhering datasets. The color of the squares in the heatmap reflects the z-score. The color reflects the direction of change for the gene expression, with red indicating positive and blue - negative z-score. Z-scores >2 indicate that the function is statistically significantly increased. Z-scores <-2 indicate that the function is statistically significantly decreased. The intensity of the colors indicates the prediction strength.

### qRT-PCR and miRNA expression analysis

Total RNA was isolated using Trizol reagent (Invitrogen/Thermo Fisher Scientific) and used for cDNA synthesis with random hexamer primers for measuring the primary transcript levels. TaqMan miRNA-specific assays (Ambion/Thermo Fisher Scientific) were used to measure miRNA-29 levels and normalized to RNU (ThermoFisher). For mRNAs, qRT-PCR was performed with SensiFast cDNA synthesis kit (Bioline) using the manufacturer’s protocol, followed by PCR using *RPL27* as reference gene.

### miRNA *in situ* hybridization combined with immunofluorescence

Based on previous work on miRNA *in situ* hybridisation analysis with fluorescently labelled LNA probes, we developed a modified miRNA-29 *in situ* hybridisation protocol where the probes specifically hybridizing miRNA-29a, miRNA-29b, or a scramble sequence of the same length are modified at the backbone and labeled with Cy2 ^6, 45^. Briefly, cryopreserved mouse wound biopsies were sectioned (14 μm) and fixed with 4% paraformaldehyde (PFA). miRNA-29a and scrambled sequences (48), synthesised and Cy2-labelled by RiboTask, were hybridised at 63° C (miRNA-29a) or 55° C (scrambled). Sections of the mouse hippocampus, which expresses high levels of miRNA-29a, were used as a positive control, whereas skin sections from *miRNA29ab1* knockout mice, hybridised with the miRNA-29a probe, as well as sections hybridised with the scrambled probe, served as negative controls to find the optimal temperature for hybridisation and washing steps ^6^. The antibody for keratin K10 (DAKO) and fibronectin (FN) were added at the last step of the wound section incubation and visualized using the secondary anti-mouse Cy3- and anti-rabbit Cy5-labelled antibody.

### Adhesion assay

Following the transfection of IFK with anti-miRNAs or single transfection with miRNA-29 mimics, the cells were seeded on fibronectin coated wells (8µg/cm^2^) and incubate for 5 minutes at 37^0^C, 5% CO2. The cells that attached within the 5 minutes were considered as fast attaching cells and the rest were slow attaching cells. After 5 minutes, the media was removed along with the slow attaching cells and fresh growth media was added. The cells were allowed to attach completely before adding PrestoBlue (Thermofischer) at 1:10 dilution to determine the number of cells that are attached. We used the 12 adhesion pathways - as per PrimePCR Disease State Panels (designed by referencing the National Library of Medicine database) to select targets for further validation.

### ECM deposition assay

DF were treated with miRNA-29 and control mimics or inhibitors and assessed for ECM proteins deposited over the course of 5 days using BCA protein assay. As fibroblasts are fast-dividing cells, the rate of dilution of the transfected oligonucleotides increases with the extended period of incubation. The ECM deposition was therefore calculated by dividing the initial transfected oligo amount by the cell number measured using PrestoBlue.

### Conditioned media from DF

HDF cells were transfected with anti-sense miRNA-29 oligonucleotides (ASO) or control ASO as mentioned before and conditioned medium (CM) was collected on the subsequent days after transfection. CM with the secreted extra cellular components was centrifuged at 2000g for 5 minutes to remove cell debris and filter sterilized using 0.22 µm filter. The medium was lyophilized in Scanvac Coolsafe Pro55’ supported by a ‘vacuubrand RC 6 hybrid’ rotary vane pump to obtain completely dried powder, which was then reconstituted using Dulbecco’s PBS to required concentration for adhesion assay with keratinocytes.

### SDS PAGE and Western blotting

Cells were harvested in 1X lysis buffer (Cell Signaling), supplemented with protease inhibitor cocktail tablets (Roche) and PhosSTOP (Roche). Cells were sonicated and left to solubilize for 30 min on ice. Then, cell lysates were centrifuged at 10,000 g for 10 min at 4°C to remove insoluble debris. Protein concentrations were measured using the BCA reagent (BioRad). Cell lysates were separated by using sodium dodecyl sulphate electrophoresis-polyacrylamide gel electrophoresis (SDS-PAGE) and transferred to nitrocellulose membranes. After blocking for 1h with Intercept blocking buffer (LI-COR) diluted 1:2 with PBS, membranes were incubated with SPARC or FERMT2 or αSMA primary antibodies diluted in antibody solution (1:2 blocking buffer/PBS containing 0.2% tween). Tubulin or GAPDH was used as a loading control. Detection was carried out with fluorescent secondary antibodies IRDye® 800CW Donkey anti-Mouse IgG and IRDye® 680CW Donkey anti-rabbit IgG visualized by LI-COR Odyssey CLx imaging system. Image Studio Analysis Software (LI-COR Bioscience) was used for imaging quantification.

### Immunofluorescence

Cells were plated onto coverslips in 12-well plates at a density of 60,000 cells per well. Next day, cells were transfected twice on subsequent days with 200 nM of anti-miRNA-29 (abc) or non-specific antisense (nsa). Then, media was changed, and cells were allowed to growth for additional 72 hours after which cells were rinsed with Dulbecco’s phosphate buffered saline (PBS) containing 0.1mM CaCl2 and 1mM MgCl2 (PBS/CM) and fixed with 2% paraformaldehyde in PBS/CM for 30 min at room temperature. After fixation, cells were washed three times with PBS/CM and permeabilized with IF buffer (PBS/CM, 0.1% Triton-X100, 0.2% BSA) for 10 min. After quenching with 50 mM NH4Cl in PBS/CM for 10 min, cells were rinsed and incubated with anti-pan collagen IV antibody or αSMA at 4°C overnight. Cells were washed with IF buffer and incubated with fluorescent secondary antibody Alexa 488 for 1 hour at room temperature. After washing, nuclei were counterstained with DAPI, and cells were then mounted using a Prolong Gold antifade medium (P10144, Invitrogen). Images were visualized using a Zeiss Axio fluorescence microscope. Fluorescence intensity was quantified in at least 10 images per condition using an in house software.

**The enzyme-linked immunosorbent assay (ELISA)** for human SPARC was performed using ready-to-go sandwitch assay kit with positive controls purchased from R&D Systems and following manifacturer’s guadlines. The standard curve was generated using recombinant SPARC, and unconditioned fibroblast medium was used as a negative control and a background. Absorbance was measured in triplicates from the conditioned medium from two independent experiments using control ASO-transfected fibroblasts and the fibroblasts transfected with miRNA-29 ASOs.

### Statistical analysis

n=3; *p<0.05, ***p<0.001, ****p<0.0001, two-way ANOVA for all data applicable unless otherwise mentioned. All experiments were done in technical and biological triplicate unless mentioned otherwise. For qRT-PCR and Luciferase assay analysis a two-tailed, paired Student’s t-test was applied (GraphPad t-test calculator).

## Author Contributions

S.K. conceived, initiated and supervised the project; L.T., R.S-A. and C.K. designed and performed experiments; Y.W., F.H., and J.H. designed the probe, modified the ASOs, and assisted in the initial miRNA-CLIP experiments; P.R. and L.T. analyzed the RNA sequencing data; H.-J.S., I.M., and P.B. developed, assisted, and analysed 3D co-cultures of human skin equivalents; L.T., R.S-A. and S.K. wrote the manuscript. All authors approved the final manuscript.

---

**Supplementary Figure S1.**
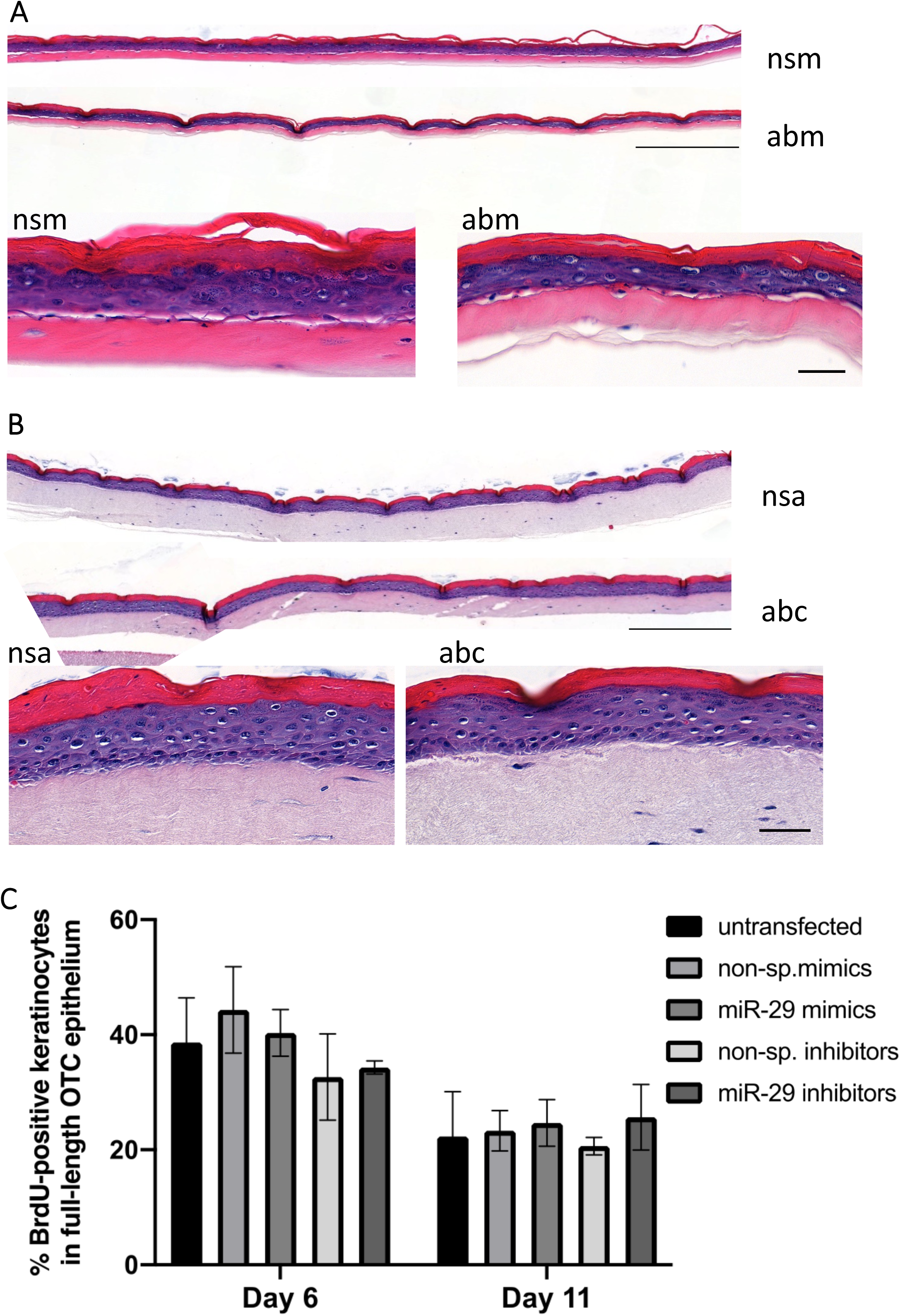
Gain of miR-29 function results in premature differentiation of human keratinocytes but does not affect their proliferation in 3D. **(A)** Human keratinocytes transfected with miR-29a/b mimics (abm) or non-specific mimics (nsm) and grown in 3D cultures at air-liquid interface for 6 days **(B)** Human keratinocytes transfected with miR-29a/b/c anti-sense oligonuclotides (abc) or non-specific oligos (nsa) and grown in 3D cultures at air-liquid interface for 6 days. Scale bar = 50μm. **(C)** Quantification of BrdU-positive keratinocytes in full-length SE epithelium expressed as % of all cells. BrdU was measured in response to transfections with non-specific mimic, non-specific inhibitor, miR-29 mimic, miR-29 inhibitor. Scale bar = 100μm.

**Supplementary Figure S2.**
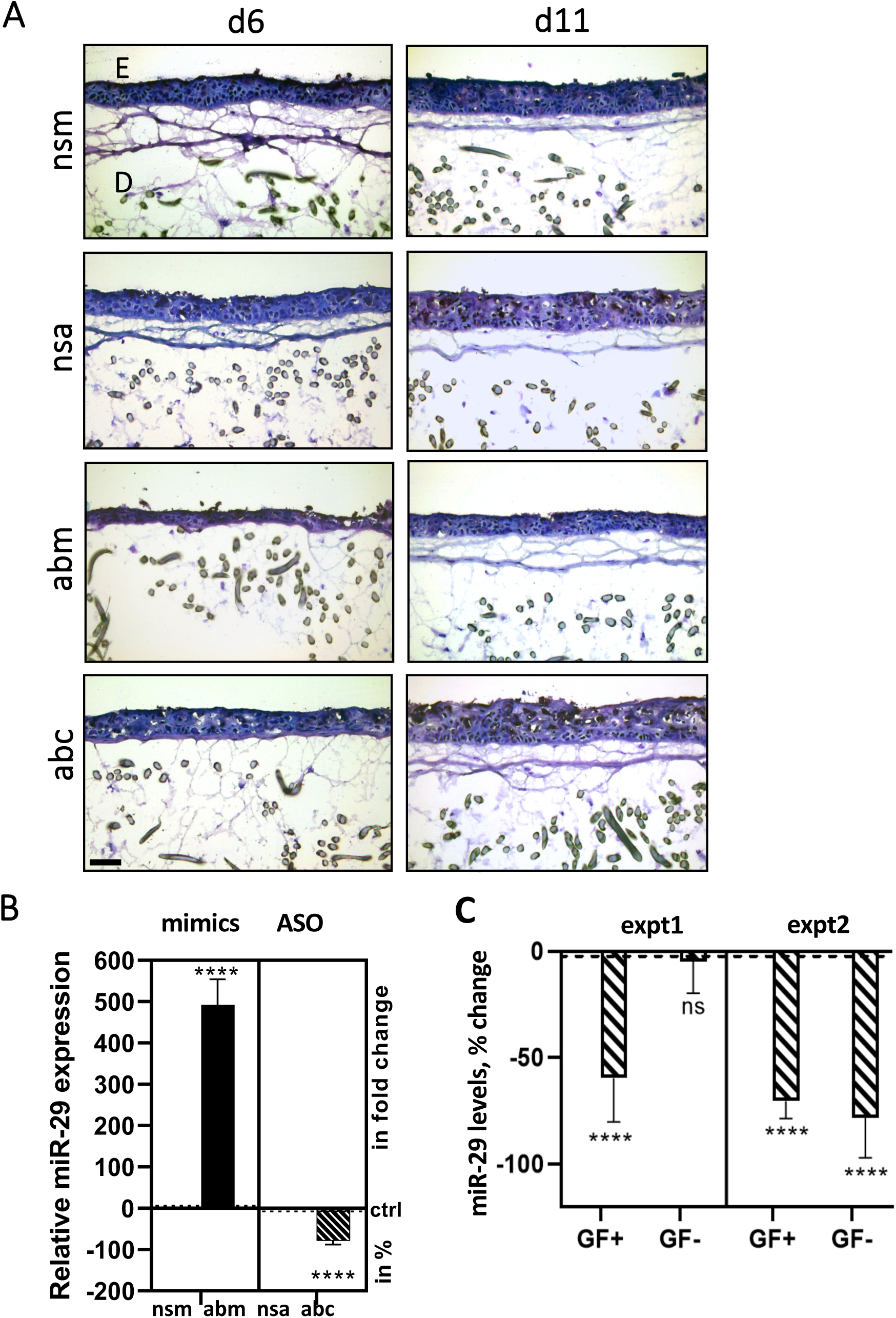
Inhibition of miR-29 permits growth and supports basal adhesion of keratinocytes. Human primary keratinocytes were double transfected with 200nM of miR-29 inhibitors (abc) or non-specific antisense (nsa) or 50nM of miR-29 mimics (abm) or non-specific sense oligo (nsm) and grown as SE. **(B)** Primary human keratinocytes were transfected with miRNA-29 mimics (abm) or inhibitors (abc) with corresponding non-specific controls (nsm and nsa). **(C)** Growth factors were withdrawn or re-added following a 24-hour starvation of transfected cells. The fast adhering cells were separated using fibronectin coated plates, total RNA was isolated and miRNA-29 levels were measured by TaqMan assays. *P < 0.05 or **P < 0.01 or \*\*\**P* < 0.001 or \*\*\*\**P* < 0.0001, two-way ANOVA followed by Šídák’s multiple comparison.

**Supplementary Figure S3.**
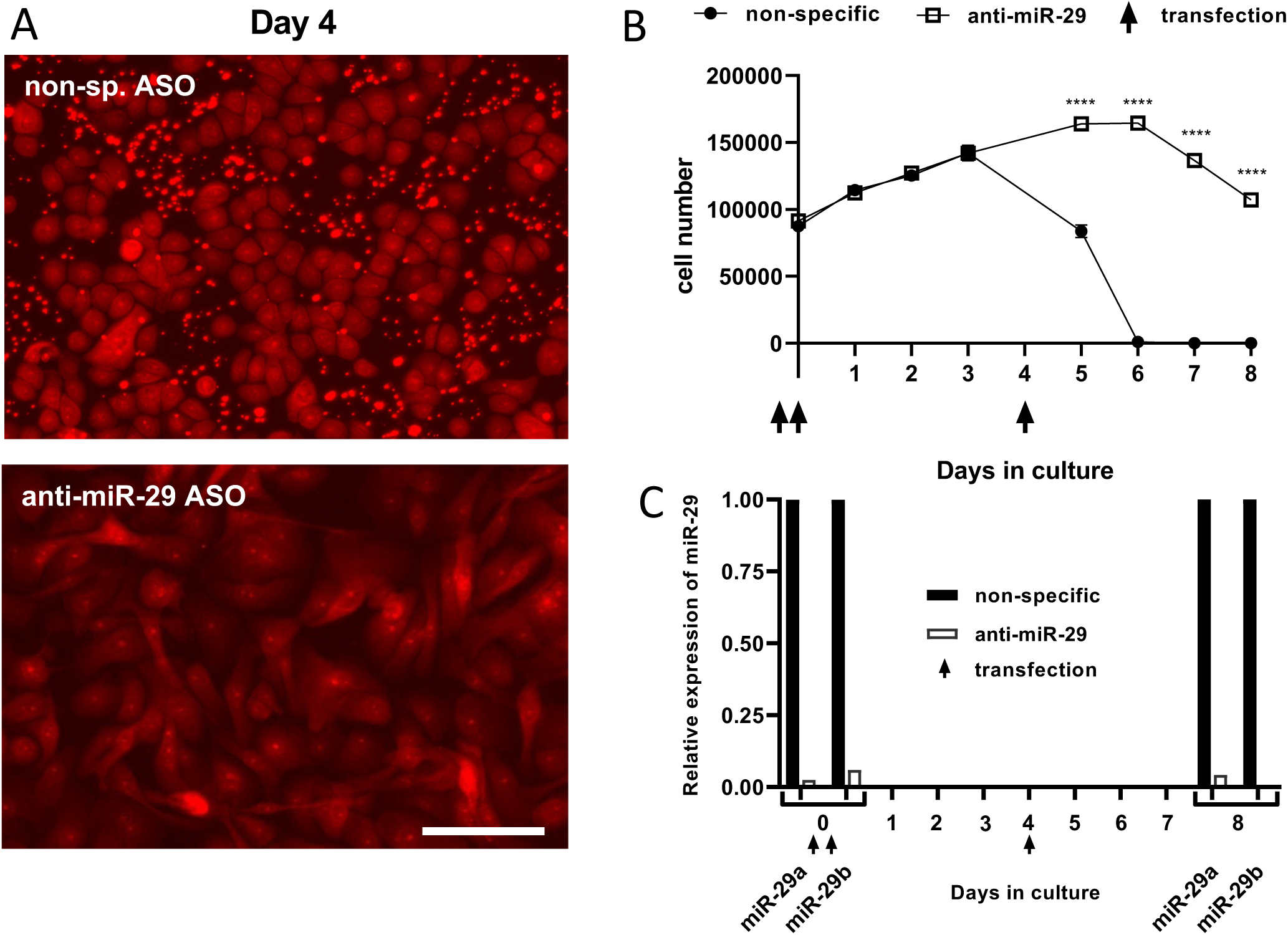
miR-29 knock-down supports keratinocyte growth. Human primary keratinocytes (P3/4) were transfected twice on subsequent days with 200nM of miR-29 inhibitors (abc) or non-specific antisense (nsa) spiked with fluorescent tagged oligos. **(A)** Representative images of the cells from each treatment after the third transfection **(B)** Cell number was determined everyday using PrestoBlue from day 1 to 8. n=3; *p<0.05, ***p<0.001, ****p<0.0001, two-way ANOVA. **(C)** Relative levels of miR-29 was measured after the second transfection to assess the knock-down/up levels and also at the end of day 8. 2-way ANOVA was performed followed by Šídák’s multiple comparison tests to ascertain the statistical significances between treatments.

**Supplementary Figure S4.**
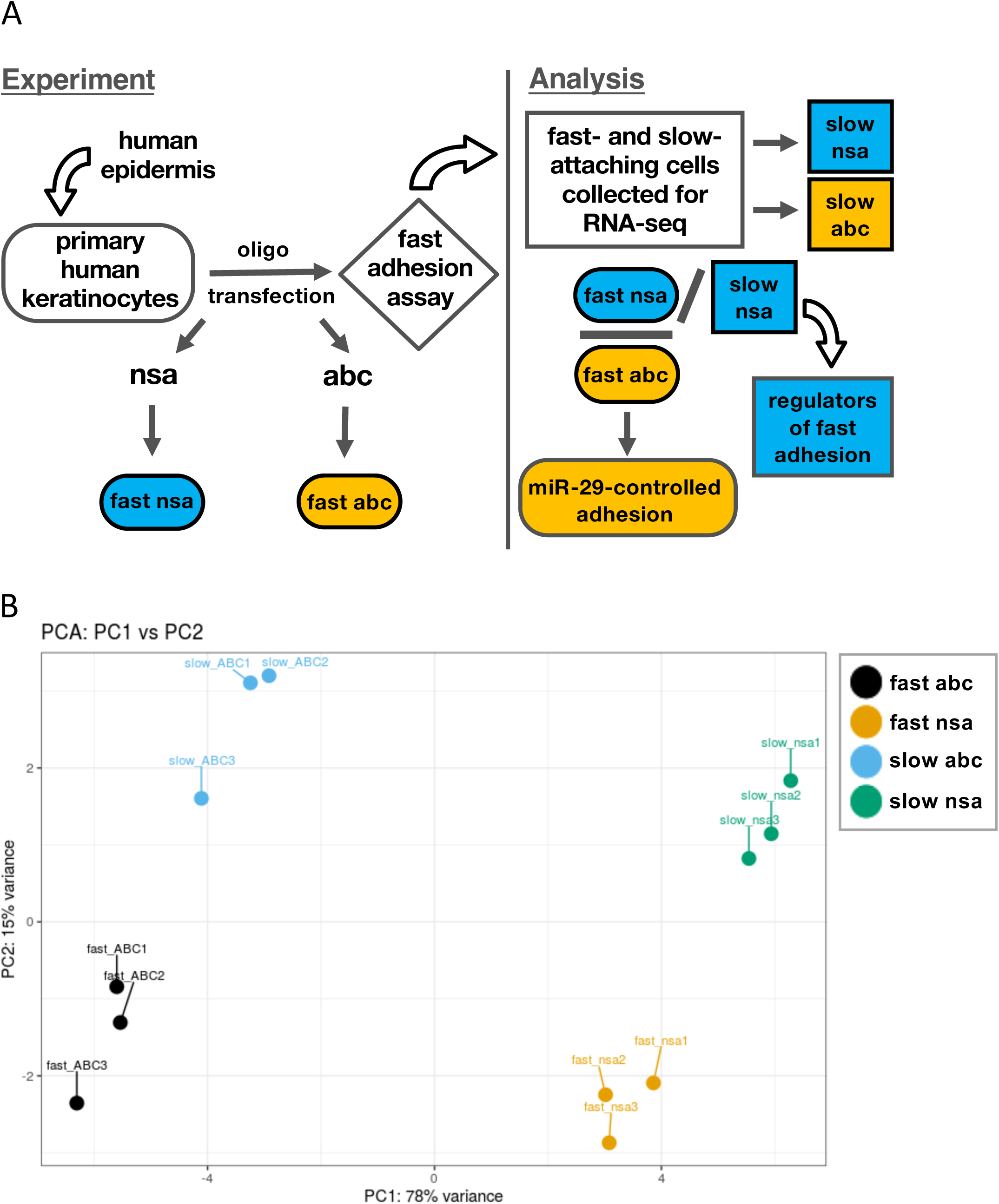
Discovering miR-29-dependent and independent mechanisms of enhanced keratinocyte adhesion. **(A)** schematic overview of experimental set-up and analysis of the mRNAs affected by knock-down of miR-29 in fast vs. slow adhering primary human keratinocytes. **(B)** Principal Component Analysis (PCA) using the rlog data for the top 500 most variable genes shows complete separation of RNAs that change expression in fast and slow adhering cells in response to transfection with the control (nsa) and miR-29 (abc) anti-sense inhibitors.

**Supplementary Figure S5.**
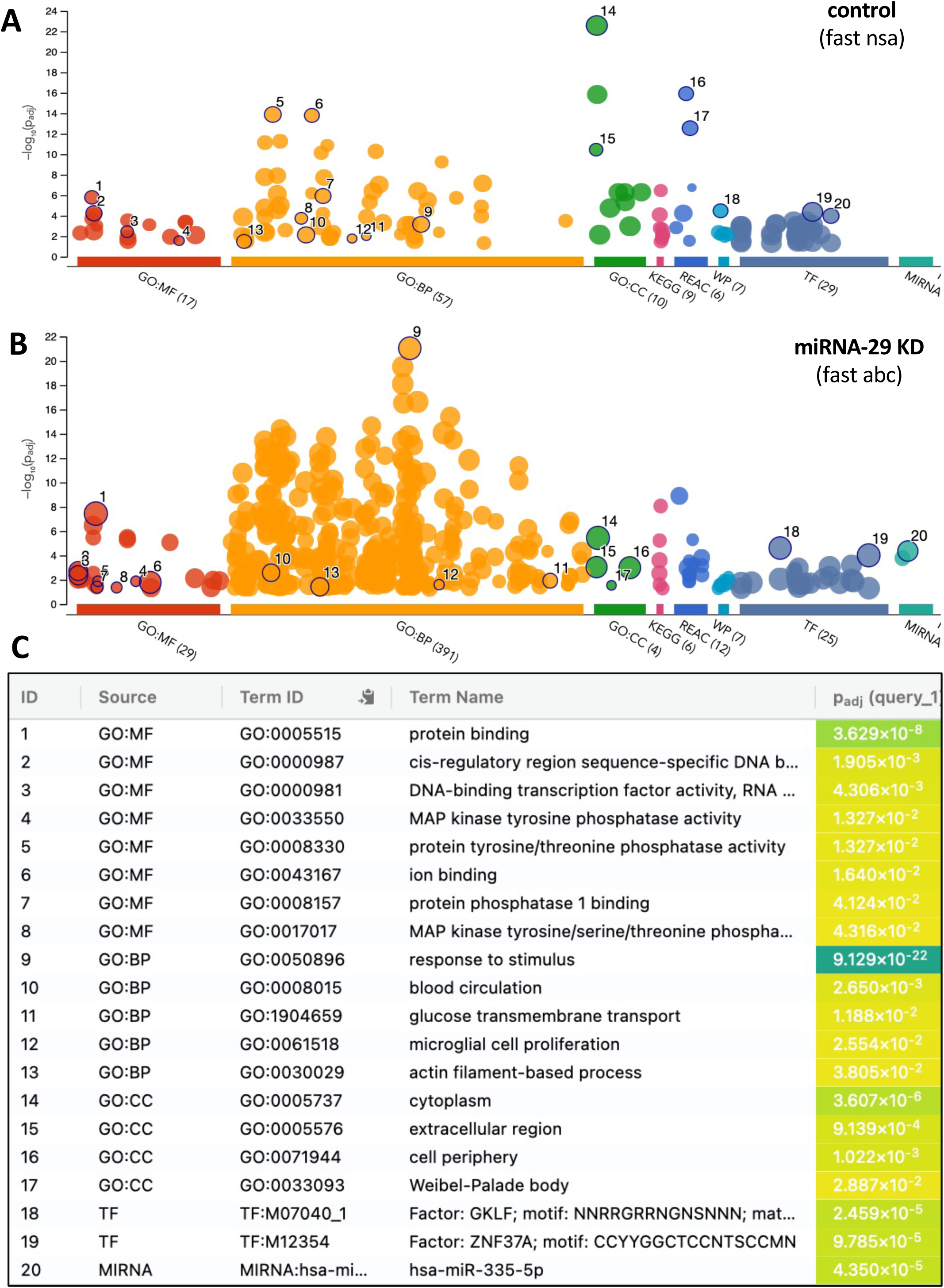
miRNA-29 regulates basal cell adhesion. Enricher-based analysis of molecular functions (MF), biological processes (BP), cellular compartments (CC), transcription factor (TF) and miRNAs upregulated upon fast adhesion in control cells **(A)** and miRNA-29 KD cells **(B). (C)** lists the results in order of significance in MF, BP, CC, TF, and miRNA categories.

**Supplementary Figure S6.**
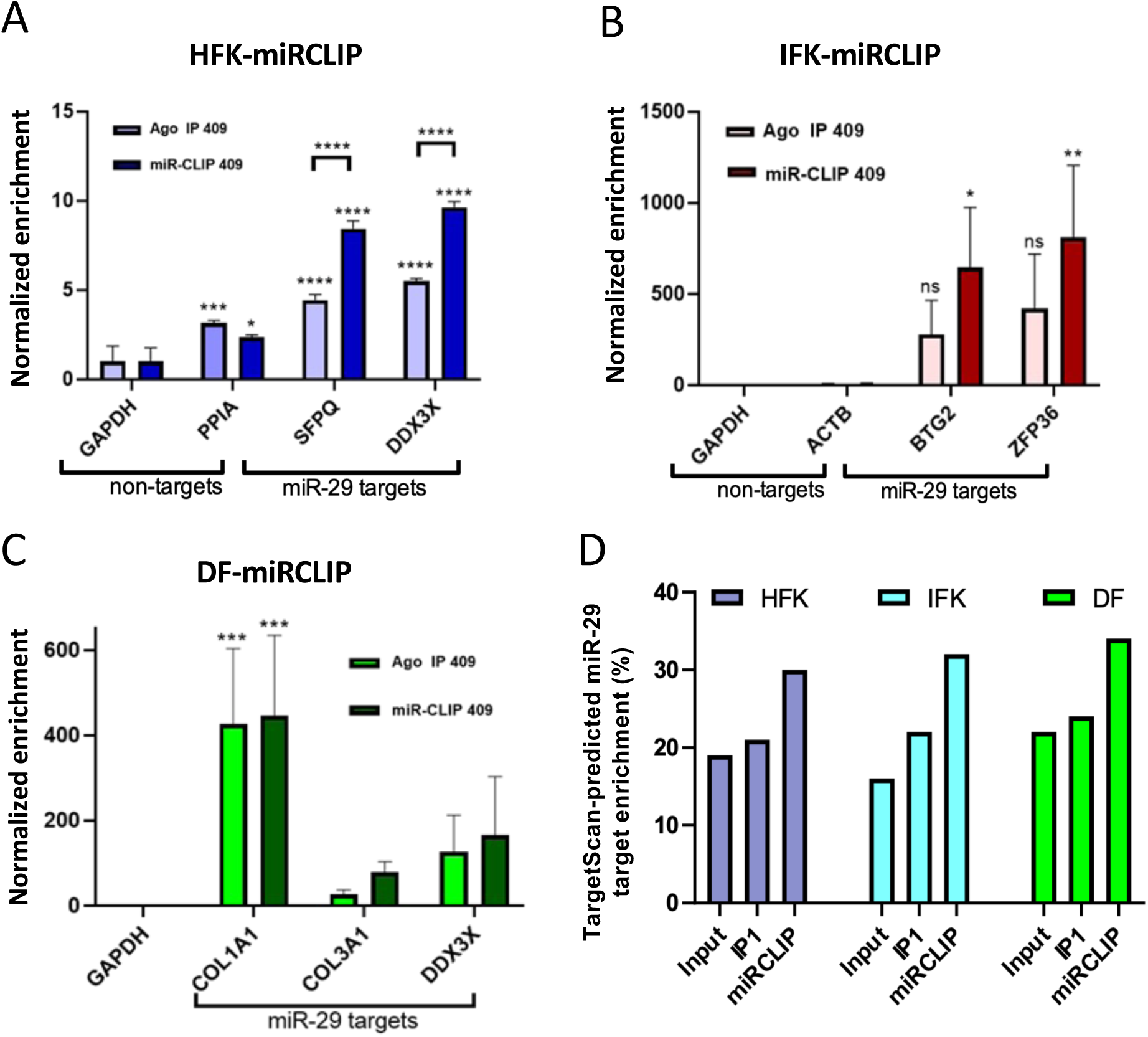
Validation of miRNA-29-CLIP in primary skin cells. Enrichment of predicted targets (as predicted in TargetScan tool) of miRNA-29 through miRNA-CLIP using the 409 probe in the first step of immunoprecipitation (IP1) and in the final step of miRNA-CLIP protocol, determined by qPCR in human follicular (HFK), interfollicular keratinocytes (IFK) and dermal fibroblasts (DF) using qPCR **(A-C)** and TargetScan **(D)** analyses.). *P < 0.05 or **P < 0.01 or \*\*\**P* < 0.001 or \*\*\*\**P* < 0.0001, two-way ANOVA, Šídák’s multiple comparisons test. Error bars indicate bidirectional standard deviation.

**Supplementary Figure S7.**
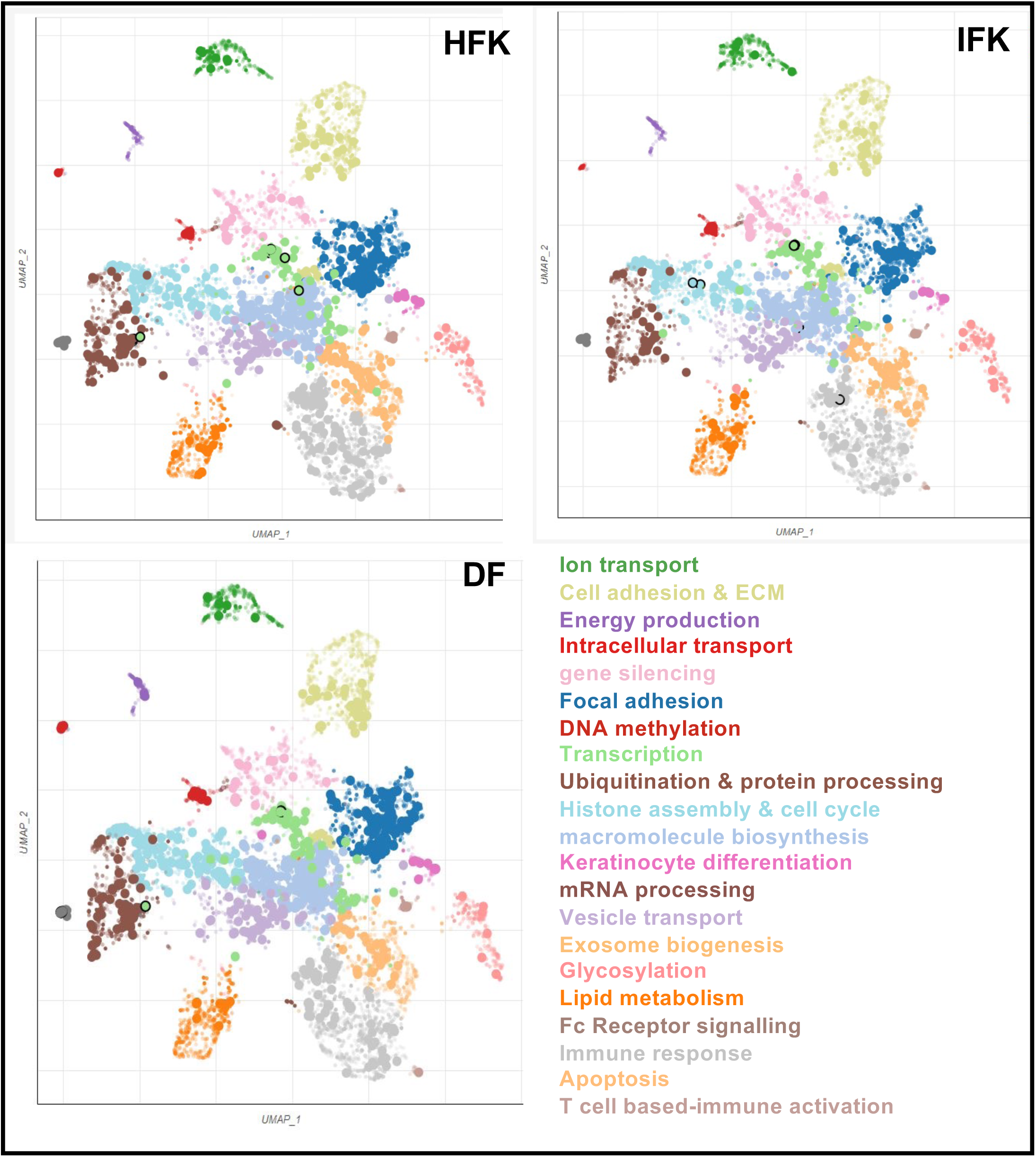
Functional analysis of miR-29-CLIP targetome in primary skin cells. Scatterplot of all terms in the GO_Biological_Processes that are represented in the miR-29 targetomes of HFK **(A),** IFK **(B)** and DF **(C)** using Enrichr Appyter. Each point represents a GO term from the targetome. The terms are plotted based on the first two Uniform Manifold Approximation and Projection (UMAP) dimensions. Terms are colored by automatically identified clusters computed with the Leiden algorithm applied to the Term frequency-inverse document frequency (TF-IDF) values. Points are plotted on the first two UMAP dimensions. The darker and larger the point, the more significantly enriched the term.

**Supplementary Figure S8.**
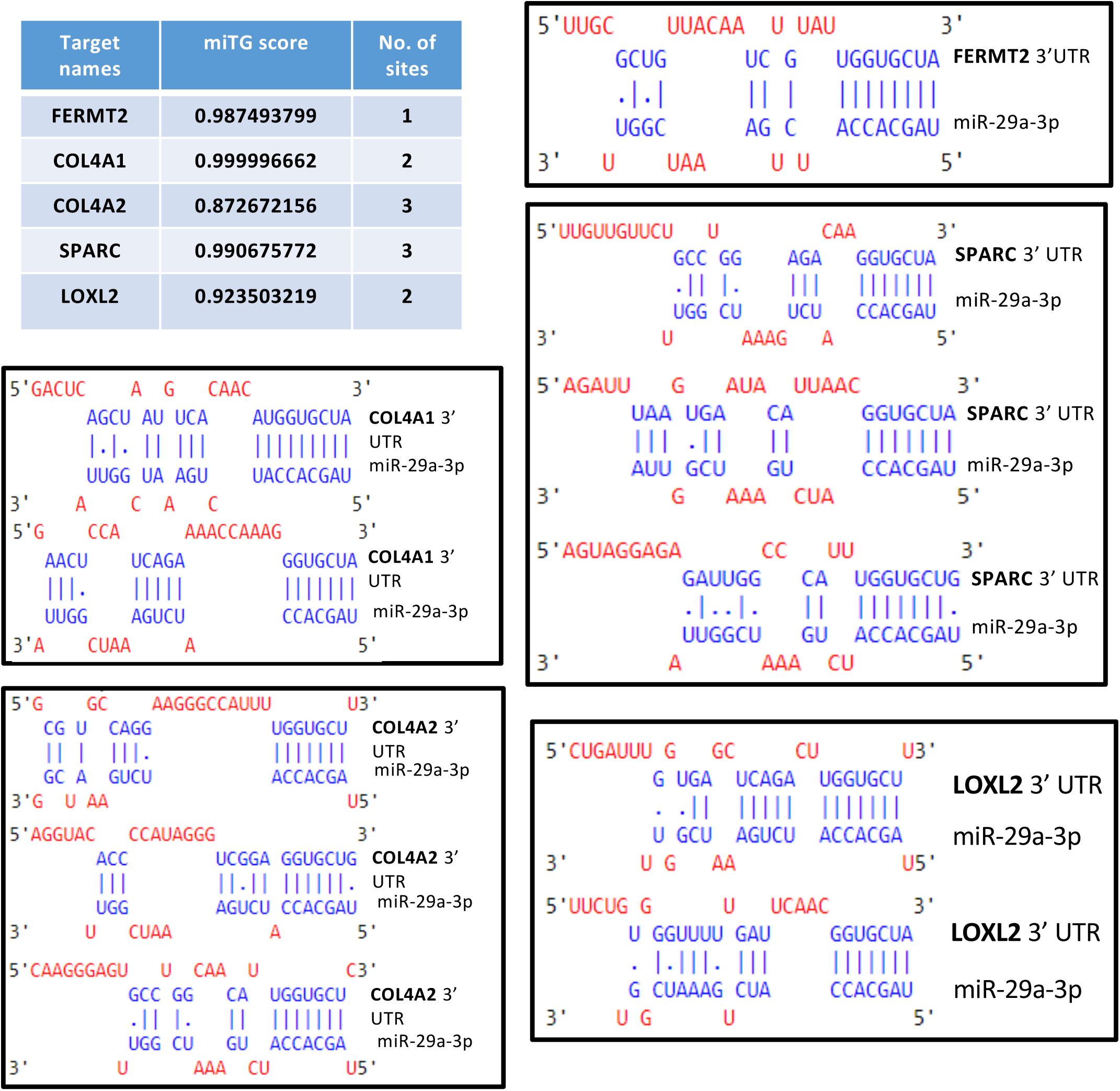
**Analysis of the ‘seed’ sequence match in miR-29 targets 3’UTRs using DIANA tool.**

**Supplementary Table 1.**
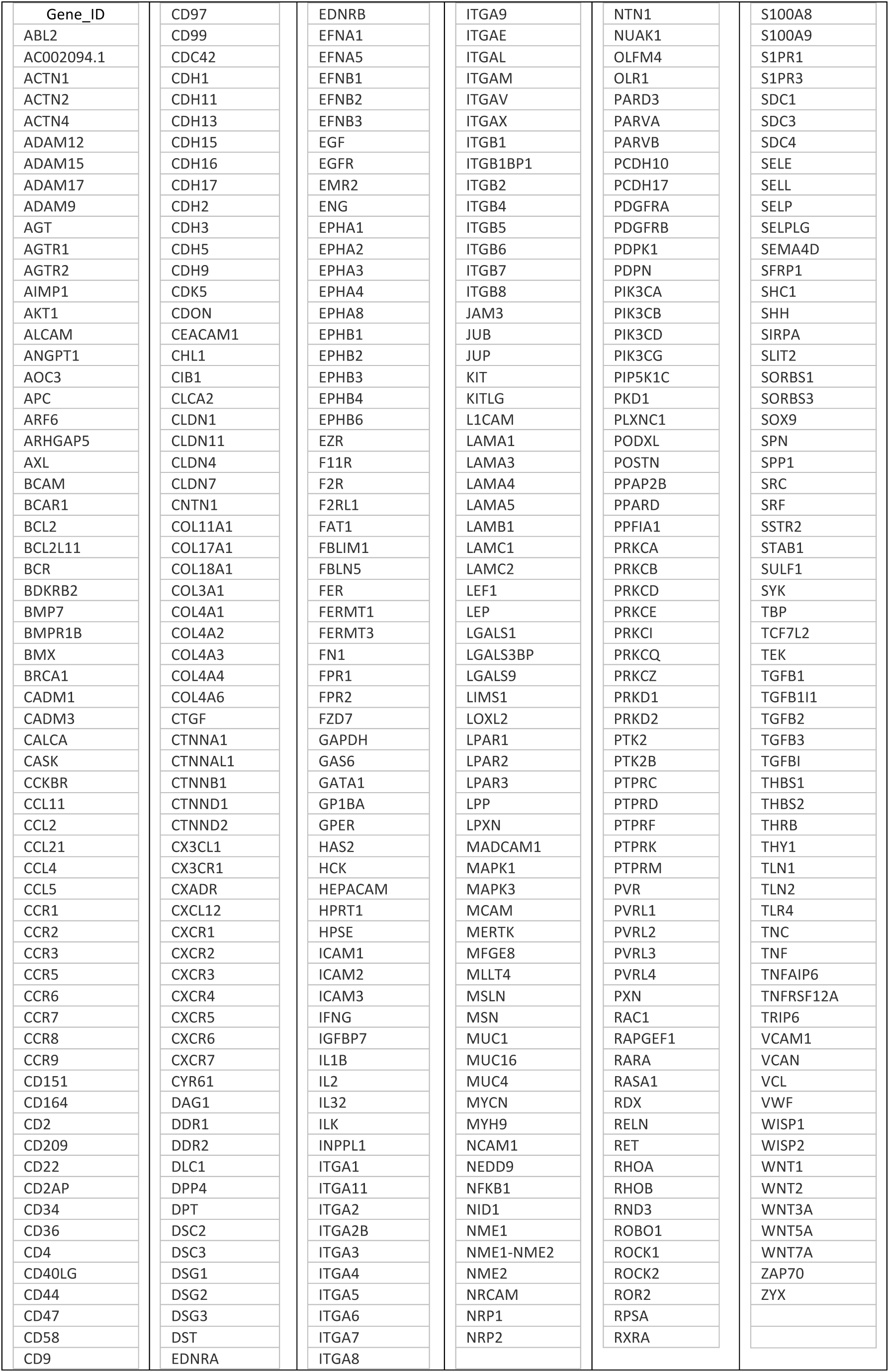
Cell adhesion pathways. Major cell adhesion pathways and the gene targets from BioRad assays.

**Supplementary Table 2.**
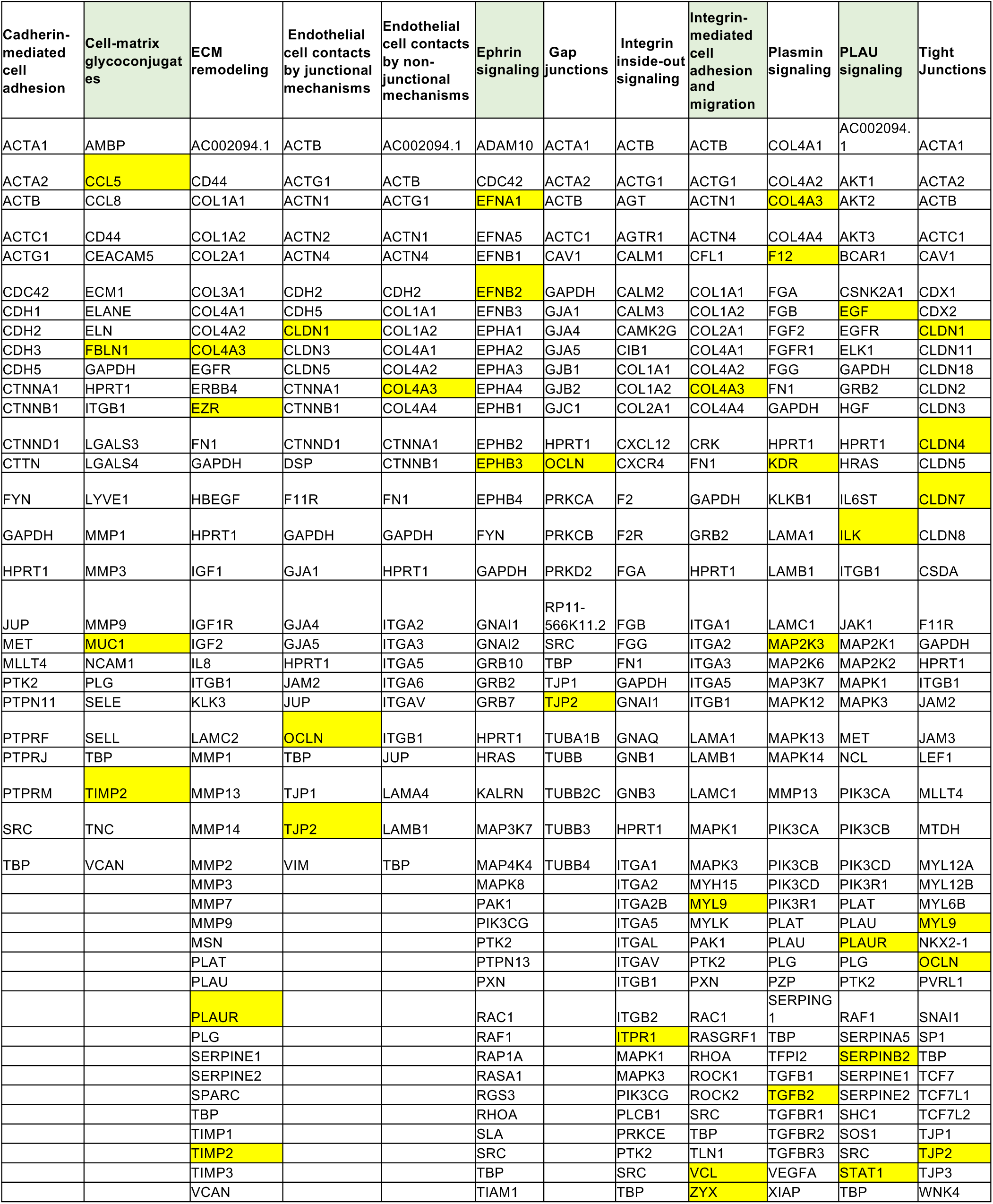
Cell adhesion pathways. Major cell adhesion pathways and the gene targets from Suppl. Table 1. Pathways that function in basal keratinocyte adhesion have green subheadings. Genes that showed significant up-regulation in the fast population of miRNA-29 LOF keratinocytes are highlighted in yellow.

**Supplementary Table 3.**
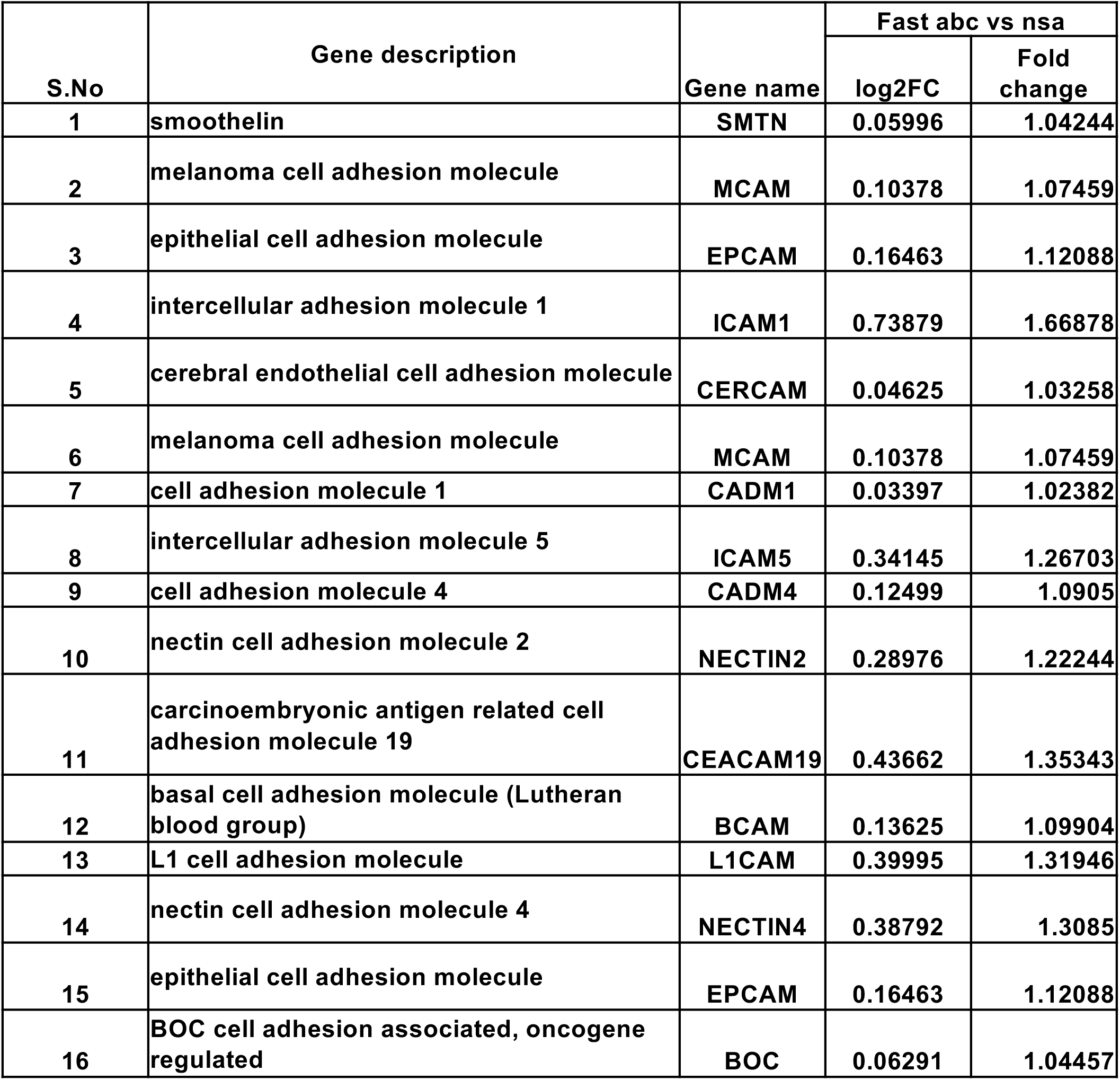
Cell Adhesion Molecules. Less studied cell adhesion molecules that showed significant up-regulation in the fast population of miRNA-29 LOF keratinocytes are highlighted in this table.

| S.No | Gene description | Gene name | Fast abc vs nsa |  |
| --- | --- | --- | --- | --- |
|  |  |  | log2FC | Fold change |
| 1 | smoothelin | SMTN | 0.05996 | 1.04244 |
| 2 | melanoma cell adhesion molecule | MCAM | 0.10378 | 1.07459 |
| 3 | epithelial cell adhesion molecule | EPCAM | 0.16463 | 1.12088 |
| 4 | intercellular adhesion molecule 1 | ICAM1 | 0.73879 | 1.66878 |
| 5 | cerebral endothelial cell adhesion molecule | CERCAM | 0.04625 | 1.03258 |
| 6 | melanoma cell adhesion molecule | MCAM | 0.10378 | 1.07459 |
| 7 | cell adhesion molecule 1 | CADM1 | 0.03397 | 1.02382 |
| 8 | intercellular adhesion molecule 5 | ICAM5 | 0.34145 | 1.26703 |
| 9 | cell adhesion molecule 4 | CADM4 | 0.12499 | 1.0905 |
| 10 | nectin cell adhesion molecule 2 | NECTIN2 | 0.28976 | 1.22244 |
| 11 | carcinoembryonic antigen related cell adhesion molecule 19 | CEACAM19 | 0.43662 | 1.35343 |
| 12 | basal cell adhesion molecule (Lutheran blood group) | BCAM | 0.13625 | 1.09904 |
| 13 | L1 cell adhesion molecule | L1CAM | 0.39995 | 1.31946 |
| 14 | nectin cell adhesion molecule 4 | NECTIN4 | 0.38792 | 1.3085 |
| 15 | epithelial cell adhesion molecule | EPCAM | 0.16463 | 1.12088 |
| 16 | BOC cell adhesion associated, oncogene regulated | BOC | 0.06291 | 1.04457 |

**Supplementary Table 4.** Oligonucleotide sequences used in analyses.

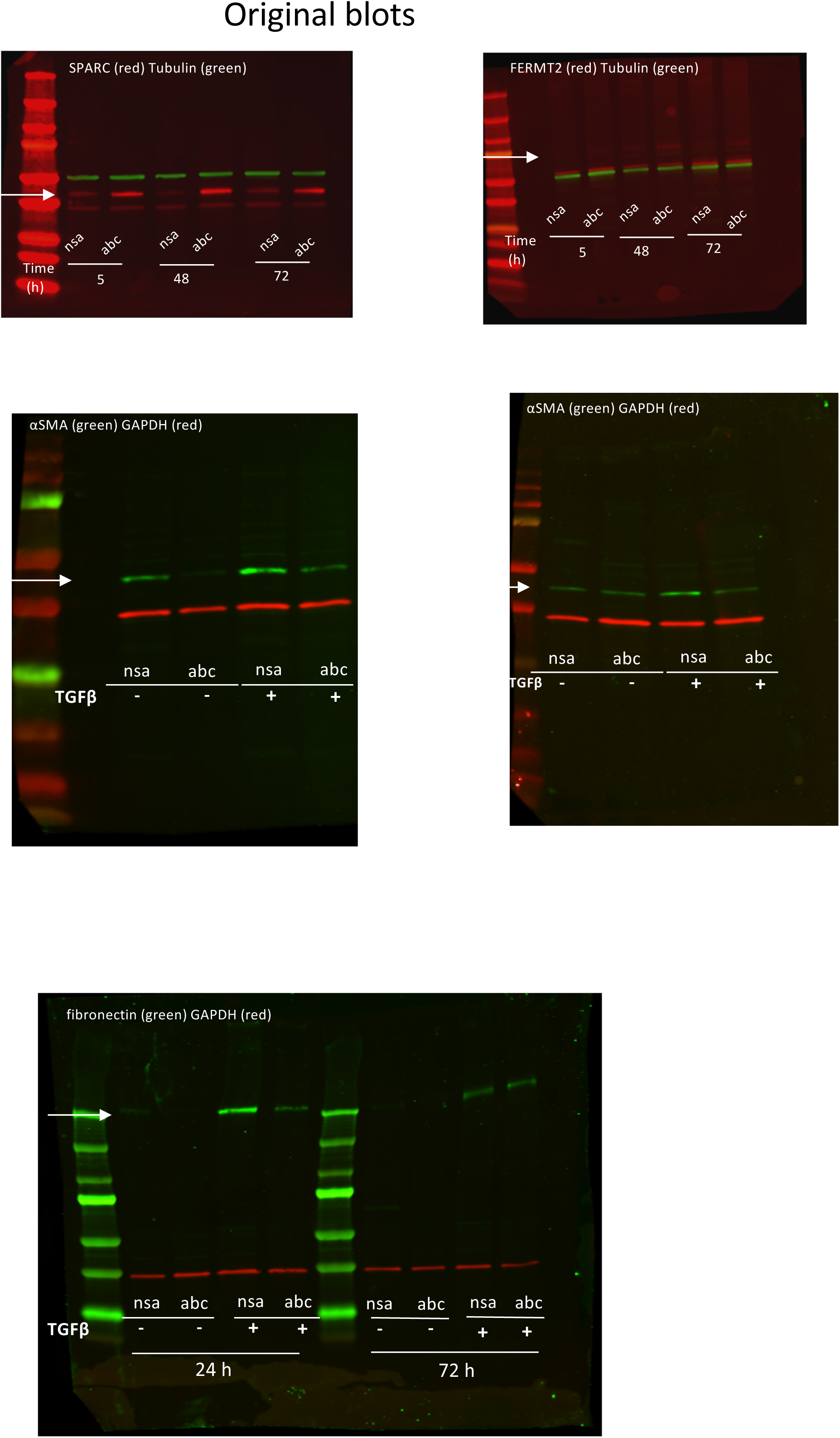

## Bibliography

1. Imig, J. et al. miR-CLIP capture of a miRNA targetome uncovers a lincRNA H19-miR-106a interaction. Nat Chem Biol 11, 107–114 (2015).

2. Bartel, D.P. Metazoan MicroRNAs. Cell 173, 20–51 (2018).

3. Wang, Y. et al. MiR-CLIP reveals iso-miR selective regulation in the miR-124 targetome. Nucleic Acids Res 49, 25–37 (2021).

4. Banerjee, J. & Sen, C.K. MicroRNAs in skin and wound healing. Methods Mol Biol 936, 343–356 (2013).

5. Hirsch, T. et al. Regeneration of the entire human epidermis using transgenic stem cells. Nature 551, 327–332 (2017).

6. Kurinna, S. et al. Interaction of the NRF2 and p63 transcription factors promotes keratinocyte proliferation in the epidermis. Nucleic Acids Res 49, 3748–3763 (2021).

7. Yang, Z. et al. MiR-29a modulates the angiogenic properties of human endothelial cells. Biochem Biophys Res Commun 434, 143–149 (2013).

8. Kurinna, S. et al. A novel Nrf2-miR-29-desmocollin-2 axis regulates desmosome function in keratinocytes. Nat Commun 5, 5099 (2014).

9. Jiang, H., Zhang, G., Wu, J.H. & Jiang, C.P. Diverse roles of miR-29 in cancer (review). Oncol Rep 31, 1509–1516 (2014).

10. Kurinna, S. & Werner, S. NRF2 and microRNAs: new but awaited relations. Biochem Soc Trans 43, 595–601 (2015).

11. Hwang, H.W., Wentzel, E.A. & Mendell, J.T. A hexanucleotide element directs microRNA nuclear import. Science 315, 97–100 (2007).

12. Fraguas, M.S. et al. MicroRNA-29 impairs the early phase of reprogramming process by targeting active DNA demethylation enzymes and Wnt signaling. Stem Cell Res 19, 21–30 (2017).

13. Gallant-Behm, C.L. et al. A MicroRNA-29 Mimic (Remlarsen) Represses Extracellular Matrix Expression and Fibroplasia in the Skin. J Invest Dermatol 139, 1073–1081 (2019).

14. Roshan, R. et al. Brain-specific knockdown of miR-29 results in neuronal cell death and ataxia in mice. RNA 20, 1287–1297 (2014).

15. 15. Yanhua Cui1, T.L., Dehua Yang, Song Li, Weidong Le miR-29 regulates Tet1 expression and contributes to early differentiation of mouse ESCs. Oncotarget 7, 64932–64941 (2016).

16. Maurer, B. et al. MicroRNA-29, a key regulator of collagen expression in systemic sclerosis. Arthritis Rheum 62, 1733–1743 (2010).

17. Eming, S.A., Martin, P. & Tomic-Canic, M. Wound repair and regeneration: mechanisms, signaling, and translation. Sci Transl Med 6, 265sr266 (2014).

18. Smola, H. et al. Dynamics of basement membrane formation by keratinocyte-fibroblast interactions in organotypic skin culture. Exp Cell Res 239, 399–410 (1998).

19. Stark, H.J. et al. Authentic fibroblast matrix in dermal equivalents normalises epidermal histogenesis and dermoepidermal junction in organotypic co-culture. Eur J Cell Biol 83, 631–645 (2004).

20. Bell, E., Ehrlich, H.P., Buttle, D.J. & Nakatsuji, T. Living tissue formed in vitro and accepted as skin-equivalent tissue of full thickness. Science 211, 1052–1054 (1981).

21. Lim, J.H., Kim, D.H., Noh, K.H., Jung, C.R. & Kang, H.M. The proliferative and multipotent epidermal progenitor cells for human skin reconstruction in vitro and in vivo. Cell Prolif 55, e13284 (2022).

22. Peterson, S.M. et al. Common features of microRNA target prediction tools. Front Genet 5, 23 (2014).

23. Paraskevopoulou, M.D. et al. DIANA-microT web server v5.0: service integration into miRNA functional analysis workflows. Nucleic Acids Res 41, W169–173 (2013).

24. Agarwal, V., Bell, G.W., Nam, J.W. & Bartel, D.P. Predicting effective microRNA target sites in mammalian mRNAs. Elife 4 (2015).

25. Horton, E.R. et al. Definition of a consensus integrin adhesome and its dynamics during adhesion complex assembly and disassembly. Nat Cell Biol 17, 1577–1587 (2015).

26. Yang, T. et al. miR-29 mediates TGFbeta1-induced extracellular matrix synthesis through activation of PI3K-AKT pathway in human lung fibroblasts. J Cell Biochem 114, 1336–1342 (2013).

27. Qin, W. et al. TGF-beta/Smad3 signaling promotes renal fibrosis by inhibiting miR-29. J Am Soc Nephrol 22, 1462–1474 (2011).

28. Knabel, M.K. et al. Systemic Delivery of scAAV8-Encoded MiR-29a Ameliorates Hepatic Fibrosis in Carbon Tetrachloride-Treated Mice. PLoS One 10, e0124411 (2015).

29. Takahashi, M., Eda, A., Fukushima, T. & Hohjoh, H. Reduction of type IV collagen by upregulated miR-29 in normal elderly mouse and klotho-deficient, senescence-model mouse. PLoS One 7, e48974 (2012).

30. Sage, E.H. Terms of attachment: SPARC and tumorigenesis. Nat Med 3, 144–146 (1997).

31. Barker, T.H. et al. SPARC regulates extracellular matrix organization through its modulation of integrin-linked kinase activity. J Biol Chem 280, 36483–36493 (2005).

32. Basu, A., Kligman, L.H., Samulewicz, S.J. & Howe, C.C. Impaired wound healing in mice deficient in a matricellular protein SPARC (osteonectin, BM-40). BMC Cell Biol 2, 15 (2001).

33. Larjava H, K.L., Häkkinen L. in In: Madame Curie Bioscience Database [Internet]. (Landes Bioscience, Austin (TX); 2000-2013).

34. Lu, J. et al. Basement Membrane Regulates Fibronectin Organization Using Sliding Focal Adhesions Driven by a Contractile Winch. Dev Cell 52, 631–646 e634 (2020).

35. Martin, P. & Nunan, R. Cellular and molecular mechanisms of repair in acute and chronic wound healing. Br J Dermatol 173, 370–378 (2015).

36. Kaur, P. & Li, A. Adhesive properties of human basal epidermal cells: an analysis of keratinocyte stem cells, transit amplifying cells, and postmitotic differentiating cells. J Invest Dermatol 114, 413–420 (2000).

37. Chen, S., Lewallen, M. & Xie, T. Adhesion in the stem cell niche: biological roles and regulation. Development 140, 255–265 (2013).

38. Hsiao, P.F. et al. Thioredoxin-interacting protein regulates keratinocyte differentiation: Implication of its role in psoriasis. FASEB J 36, e22313 (2022).

39. Roca-Cusachs, P. et al. Integrin-dependent force transmission to the extracellular matrix by alpha-actinin triggers adhesion maturation. Proc Natl Acad Sci U S A 110, E1361–1370 (2013).

40. Luna, C., Li, G., Qiu, J., Epstein, D.L. & Gonzalez, P. Cross-talk between miR-29 and transforming growth factor-betas in trabecular meshwork cells. Invest Ophthalmol Vis Sci 52, 3567–3572 (2011).

41. Kim, B.N. et al. TGF-beta induced EMT and stemness characteristics are associated with epigenetic regulation in lung cancer. Sci Rep 10, 10597 (2020).

42. Song, H., Ding, L., Zhang, S. & Wang, W. MiR-29 family members interact with SPARC to regulate glucose metabolism. Biochem Biophys Res Commun 497, 667–674 (2018).

43. Qiu, F. et al. miR-29a/b enhances cell migration and invasion in nasopharyngeal carcinoma progression by regulating SPARC and COL3A1 gene expression. PLoS One 10, e0120969 (2015).

44. Nishikawa, R. et al. Tumour-suppressive microRNA-29s directly regulate LOXL2 expression and inhibit cancer cell migration and invasion in renal cell carcinoma. FEBS Lett 589, 2136–2145 (2015).

45. Dooley, J. et al. The microRNA-29 Family Dictates the Balance Between Homeostatic and Pathological Glucose Handling in Diabetes and Obesity. Diabetes 65, 53–61 (2016).

